# Binding of phosphatidylserine-positive microparticles by PBMCs classifies disease severity in COVID-19 patients

**DOI:** 10.1101/2021.06.18.448935

**Authors:** Lisa Rausch, Konstantin Lutz, Martina Schifferer, Elena Winheim, Rudi Gruber, Linus Rinke, Johannes C. Hellmuth, Clemens Scherer, Maximilian Muenchhoff, Christopher Mandel, Michael Bergwelt-Baildon, Mikael Simons, Tobias Straub, Anne B. Krug, Jan Kranich, Thomas Brocker

**Affiliations:** Institute for Immunology, Biomedical Center (BMC), Faculty of Medicine, LMU Munich, Munich, Germany; German Center for Neurodegenerative Diseases (DZNE), Munich, Germany; Munich Cluster of Systems Neurology (Synergy), Munich, Germany; bene pharmaChem GmbH & Co.KG. Geretsried, Germany; Department of Medicine III, University Hospital, LMU Munich, Munich, Germany; COVID-19 Registry of the LMU Munich (CORKUM), University Hospital, LMU Munich, Munich, Germany; Department of Medicine I, University Hospital, LMU Munich, Munich, Germany; Max von Pettenkofer Institute & Gene Center, Virology, National Reference Center for Retroviruses, LMU München, Munich, Germany; German Center for Infection Research (DZIF), partner site Munich, Germany; Department of Medicine IV, University Hospital, LMU Munich, Munich, Germany; Institute of Neuronal Cell Biology, Technical University of Munich, Munich, Germany; Core facility Bioinformatics, Biomedical Center (BMC), Faculty of Medicine, LMU Munich, Munich, Germany

**Keywords:** Thromboinflammation, lymphopenia, SARS-CoV-2, COVID-19, phosphatidylserine, apoptosis, platelet-derived microparticle, CD8^+^ T cells

## Abstract

Infection with SARS-CoV-2 is associated with thromboinflammation, involving thrombotic and inflammatory responses, in many COVID-19 patients. In addition, immune dysfunction occurs in patients characterized by T cell exhaustion and severe lymphopenia. We investigated the distribution of phosphatidylserine (PS), a marker of dying cells, activated platelets, and platelet-derived microparticles (PMP), during the clinical course of COVID-19. We found an unexpectedly high amount of blood cells loaded with PS^+^ PMPs for weeks after the initial COVID-19 diagnosis. Elevated frequencies of PS^+^PMP^+^ PBMCs correlated strongly with increasing disease severity. As a marker, PS outperformed established laboratory markers for inflammation, leucocyte composition, and coagulation, currently used for COVID-19 clinical outcome prognosis. PS^+^ PMPs preferentially bound to CD8^+^ T cells with gene expression signatures of proliferating effector rather than memory T cells. As PS^+^ PMPs carried programmed death-ligand 1 (PD-L1), they may affect T cell expansion or function. Our data provide a novel marker for disease severity and show that PS, which can trigger the blood coagulation cascade, the complement system, and inflammation, resides on activated immune cells. Therefore, PS may serve as a beacon to attract thromboinflammatory processes toward lymphocytes and cause immune dysfunction in COVID-19.

## INTRODUCTION

The recently emerged human pathogenic severe acute respiratory syndrome coronavirus 2 (SARS-CoV-2) causes various clinical syndromes, summarized under coronavirus disease 2019 (COVID-19). The excessive inflammatory response associated with COVID-19 can cause severe complications such as acute respiratory distress syndrome, septic shock, and multi-organ failure ^1-3^. A significant cause of morbidity and mortality in COVID-19 patients is ‘thromboinflammation’. Although not fully understood, inflammation through complement activation and cytokine release, platelet overactivity and apoptosis (thrombocytopathy), as well as coagulation abnormalities (coagulopathy) play critical roles in this complex clinical picture (reviewed in ^4^).

Similar vascular complications occur in the antiphospholipid syndrome, where autoantibodies target phosphatidylserine (PS) / prothrombin complexes ^5^. This syndrome may manifest itself in COVID-19 patients ^6^, linking PS to thromboinflammation. PS is a plasma membrane component actively retained by an ATP-requiring process at the inner membrane surface in living cells. PS retention stops, for example, during cell death or when cells release PS-containing microparticles or enveloped viruses. Then PS relocates to the outer layer of the cell membrane, where it can interact with extracellular proteins, including coagulation and complement systems. PS activates the alternative and the classical complement pathways ^7-9^ by binding to complement C3 ^10^, C3bi ^8^, and C1q ^11^. Activated platelets release platelet-derived microparticles (PMPs), which cause thrombin formation, coagulation, activation of the complement system, and inflammation in a PS-dependent manner ^12-14^.

Patients with severe COVID-19 also show striking immune dysregulation, the reasons for which are not entirely understood. A direct correlation between blood clotting components and the immune response exists ^15^. Various immune abnormalities such as increased inflammatory cytokines ^16^, immune cell exhaustion ^17^, and general lymphopenia ^18-22^ correlate with disease severity ^23^. T cell lymphopenia ^24^, probably caused by excessive apoptotic T cell death similar to sepsis ^25^, is of particular relevance as SARS-CoV-2-specific T cell responses control and resolve the primary infection ^26-29^. In fatal COVID-19, the adaptive immune response starts too late (reviewed in ^30^), while its rapid onset would be highly beneficial ^27,31,32^. However, the reasons and precise mechanisms for adaptive immune disturbance, lymphopenia, and thromboinflammation in COVID-19 remain poorly defined.

To investigate these aspects of COVID-19 in more detail, we interrogated PBMC of 54 patients from the COVID-19 Registry of the LMU Munich (CORKUM) and 35 healthy and 12 recovered donors between April 2020 and February 2021. We performed image flow cytometry (IFC) and image analysis by deep learning algorithms ^33^ using highly sensitive reagents specific for PS ^33,34^. COVID-19 blood samples contained abnormally high numbers of PS^+^ peripheral blood mononuclear cells (PBMC). Although PS is a marker for dying cells, nearly all PS^+^ cells were living cells associated with PS^+^CD41^+^ PMPs or larger PS^+^CD41^+^ platelet fragments. The grade of PS^+^ PMP-associated PBMC correlated with lymphopenia and disease severity, showing a higher correlation than commonly used laboratory diagnostic markers such as IL-6, D-Dimer ^23^, and C-reactive protein (CRP) ^35^. PS^+^ PMPs were strongly associated with dividing effector CD8^+^ T cells with upregulated expression of cell-cycle genes. Fractions of T cell-associated PS^+^ PMPs carried CD274 (PD-L1), which could impact the survival of T cells and potentially contribute to functional inhibition and lymphopenia. As PS^+^ PMPs remained associated with PBMC several weeks after the initial SARS-CoV2-diagnosis, they might sustain the adverse inflammatory and prothrombotic effects over a long time and contribute to the complex clinical picture of thromboinflammation (reviewed in ^4,36^). Together, our findings reveal an extensive association of PS^+^ PMPs with lymphocytes as a novel marker to classify COVID-19 disease severity and a potentially relevant contributor to thromboinflammation and lymphocyte dysfunction.

## RESULTS

### Lymphocytes from COVID-19 patients show substantial PS surface exposure

To test if immune cell death contributes to lymphopenia during COVID-19, we analyzed peripheral blood mononuclear cells (PBMC) of COVID-19 patients and compared them to those from healthy and recovered donors. Suppl. Tables 1-3 show clinical metadata for our cohort of COVID-19 patients and control groups. One hallmark of apoptotic cell death is the PS exposure on the outer membrane surface of cells. To reveal PS on PBMC, we utilized recombinant Milk fat globule-EGF factor 8 protein (MFG-E8) derived recombinant proteins, which bind PS under physiological conditions with high sensitivity on apoptotic cells and subcellular PS^+^ extracellular vesicles (EVs) ^33,34^. Flow cytometry results showed that the frequencies of PS^+^ cells in blood from all COVID-19 patients were significantly higher than in PBMC from healthy or recovered donors (Fig. 1A). To analyze if this data can classify patients according to disease severity, we employed the World Health Organization’s (WHO) eight-point ordinal scale for COVID-19 trial endpoints ^37^ (Fig. 1B). In our patient cohort, the scores WHO 2 and WHO 7 were absent. We combined WHO scores into ‘mild’ (WHO 1–3), ‘moderate’ (WHO 4), and ‘severe’ (WHO 5–8) groups for the subsequent analyses. Additionally, we also included a group of healthy donors (HD, n = 30) and recovered patients (n = 12, >69 days post 1^st^ SARS-CoV-2^+^ diagnosis by PCR, either never hospitalized or released from the hospital with WHO score 1 – 2). The frequencies of PS^+^ PBMC increased with severity of COVID-19 disease in the following order: healthy controls (WHO 0) < recovered patients < WHO 1-3 (mild) < WHO 4 (moderate) < WHO 5-8 (severe) (Fig. 1C). In severely diseased patients, 30 – 90% of all PBMC were PS^+^ (Fig. 1C). Accordingly, the individual WHO scores positively correlated with high significance with the frequencies of PS^+^ PBMC of COVID-19 patients (Fig. 1D).

**Figure 1:**
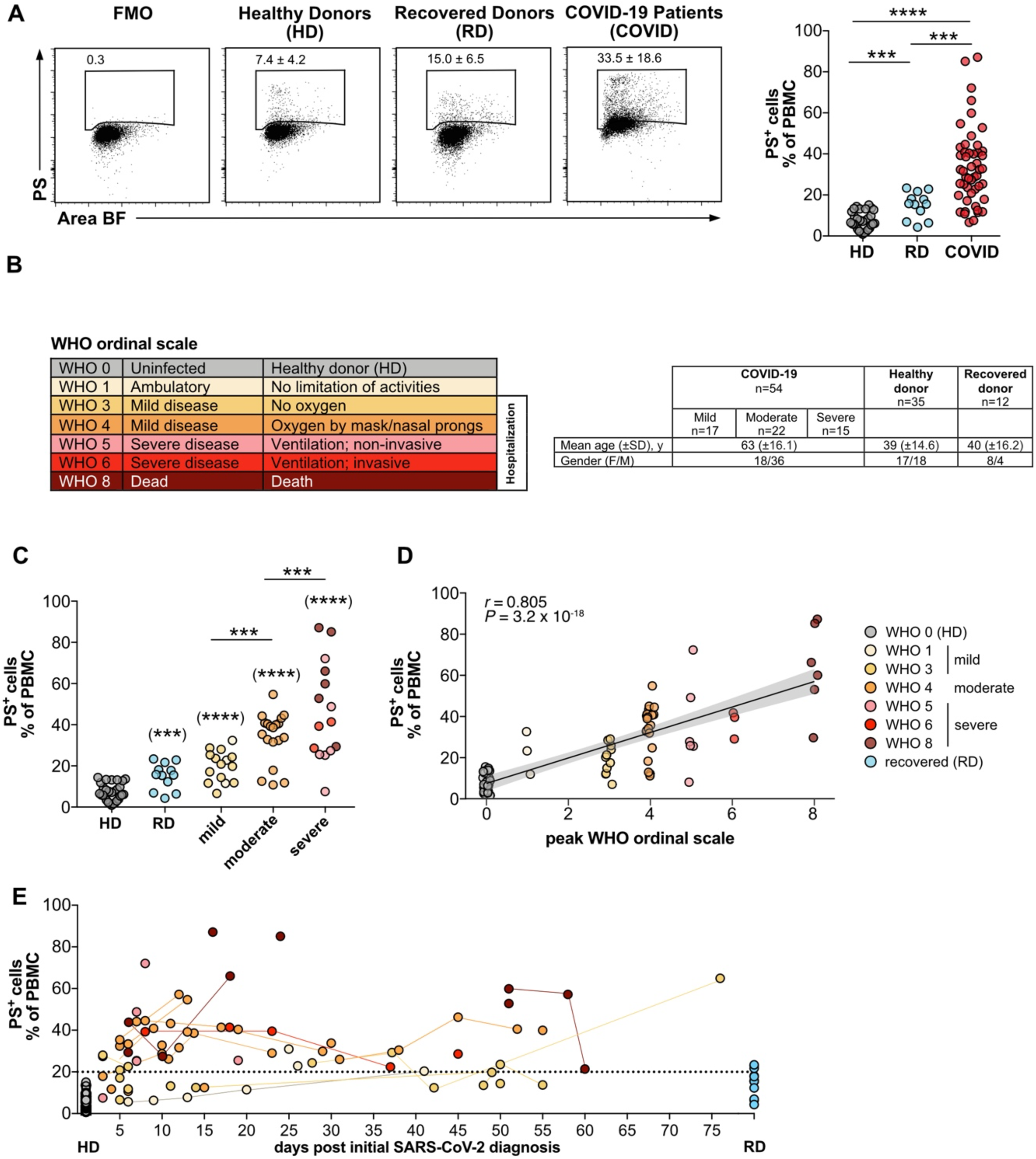
The frequencies of PS^+^ PBMC from COVID patients correlates with disease severity. **(A)** PBMC from COVID-19 patients (COVID), healthy donors (HD), and recovered donors (RD) were stained for PS and analyzed by flow cytometry. Numbers in dot plots correspond to the percentage of PS^+^ cells in the gate shown. The right-hand graph shows the summary of all percentages. **(B)** Overview of WHO ordinal scale and color code used. Table shows the number, age and gender of the different study groups. **(C)** Grouped analysis of the data from (A). **(D)** Same as (C), but plotted against the WHO ordinal scale (n = 38-79) (B). PS^+^ PBMCs correlate with the severity of the disease. The plot shows the Spearman correlation test and linear regression line with 95% confidence interval shading (A, C, D: HD, n = 30; RD, n = 12; COVID, n = 49). **(E)** Analysis of PS^+^ PBMC as shown in (A) plotted against days after initial SARS-CoV-2 diagnosis. Lines connect the same donors. Significance was determined by Mann-Whitney test: *P < 0.05, **P < 0.01, ***P < 0.001, and ****P < 0.0001. Asterisks in brackets show statistically significant differences as compared to HD.

Moreover, when we analyzed the frequencies of PS^+^ PBMC over time, we found elevated levels for up to 10 weeks in some patients (Fig. 1E), indicating that the presence of PS^+^ PBMC in COVID-19 patients might be long-lasting. However, in recovering patients, PS^+^ PBMC returned to the levels of healthy controls (Fig. 1E). In summary, the number of PS^+^ PBMC in the blood of COVID-19 patients represents a new parameter that correlates with strongly with disease severity.

### PBMC in COVID-19 patients are associated with PS^+^ EVs, and the amount correlates with the severity of the disease

PS exposure occurs on various cells and cell-derived microparticles, including tumor cells, erythrocytes, neutrophils, monocytes, endothelial cells, activated platelets, and PMPs. PS-exposure is a significant regulator of the blood coagulation system (reviewed in ^38^). We have recently shown that most PS^+^ cells in the spleen of virus-infected mice are not apoptotic, but cells carried PS^+^ extracellular vesicles ^33^. To determine whether PS^+^ PBMC in COVID-19 patients were apoptotic cells contributing to the described lymphopenia or EV^+^ cells, we analyzed the images of PS^+^ PBMC acquired by IFC. Some cells showed almost entirely PS^+^ cell bodies with strongly labeled apoptotic blebs, typical for cell death by apoptosis (Fig. 2A). These cells still have an intact cell membrane since they did not stain with the live/dead dye used to exclude necrotic cells from the analysis. However, we also detected many cells with the characteristic round brightfield image morphology of living cells, with only one or a few intensely PS^+^ structures of subcellular size (Fig. 2A). These particles resembled cell-associated PS^+^ EVs, which we recently identified in virus-infected mice ^33^. We next used a machine learning-based convolutional autoencoder (CAE) to group PBMC into *bona fide* PS^+^ apoptotic cells or cells associated with PS^+^ EVs (Fig. 2A) ^33^. After training with pre-defined images of PS^+^ dying or PS^+^ EV-associated cells, the CAE algorithm digitally sorts PS^+^ cells into both categories with high precision (Fig. 2A). When we analyzed the CAE-classified PS^+^ dying cells and PS^+^EV^+^ cells separately (Fig. 2A), we found that the majority of PS^+^ PBMC contained live cells associated with PS^+^ EV-like structures rather than PS^+^ dying cells (Fig. 2B). Despite the rarity of dying cells, patients with the score WHO 4 showed the highest frequencies (Fig. 2B, upper panel). Furthermore, only PS^+^EV^+^ PBMC, not PS^+^ dying cells, classified the patients into disease score groups mild’ (WHO 1–3), ‘moderate’ (WHO 4), and ‘severe’ (WHO 5–8) and separated them clearly from HD and recovered patients (Fig. 2B, lower panel). Similarly, PS^+^EV^+^ PBMC, but not PS^+^ dying cells, showed a highly significant correlation with WHO scores (Fig. 2C). Several laboratory values that are either increased (leukocytes, IL-6, neutrophils, procalcitonin (PCT), C-reactive protein (CRP), partial thromboplastin time (PTT), D-dimer, etc.) or decreased (lymphocytes) were shown to indicate an unfavorable progression of COVID-19 disease ^39^.

**Fig. 2:**
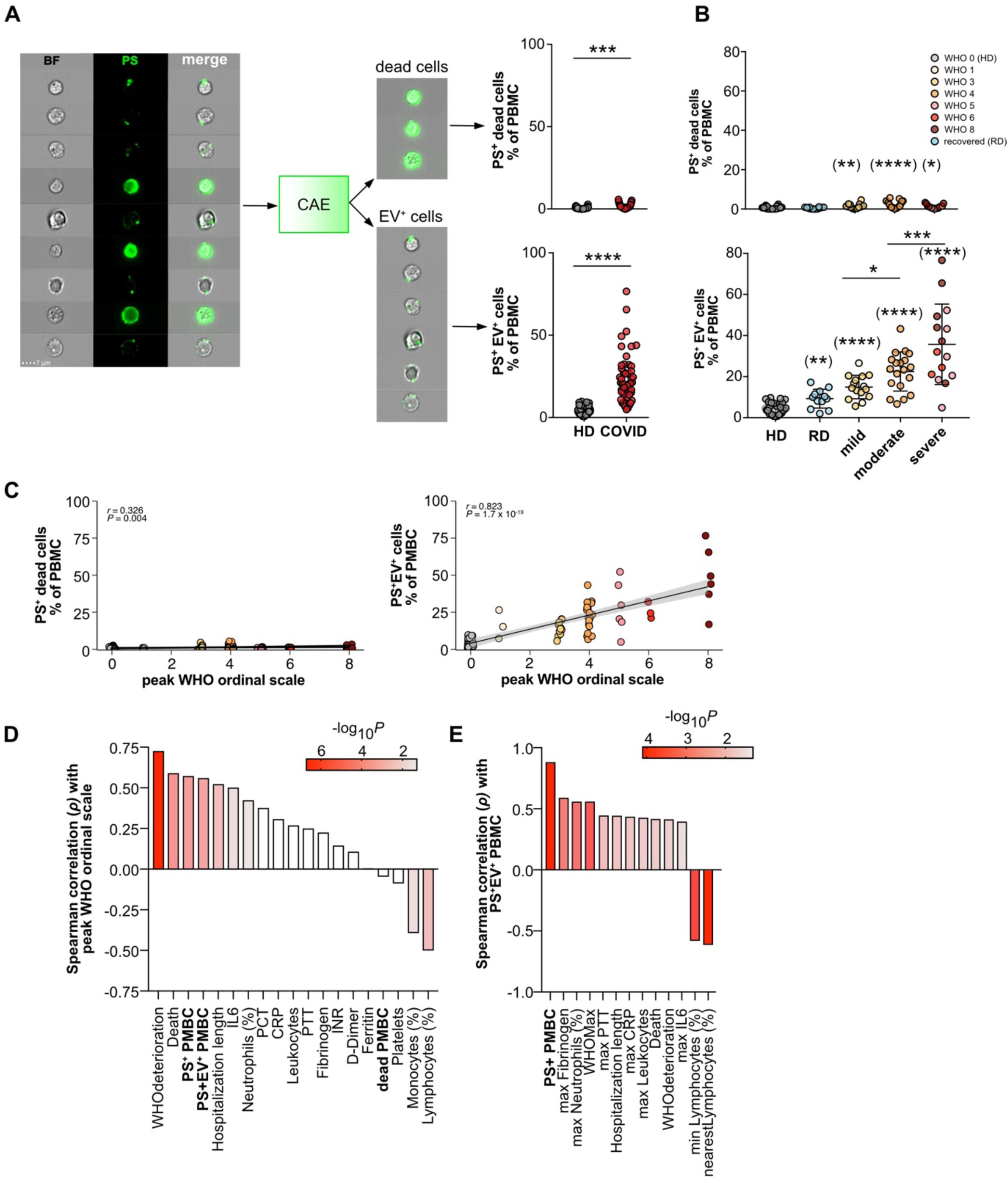
Deep learning discriminates apoptotic and EV^+^ cells. **(A)** PBMC from healthy donors (HD, n = 30) and COVID (n = 49) were analyzed by imaging flow cytometry (IFC). To discriminate apoptotic and EV^+^ cells, PBMC were analyzed using IDEAS, CAE, and FlowJo. PS^+^ cells were gated, and their TIF images (16-bit, raw) exported using the IDEAS software. CAE results with the classification apoptotic/EV^+^ were re-imported into IDEAS, and separate FCS files containing all cells or only PS^+^/dying cells and PS^+^/EV^+^ cells were generated for further analysis in FlowJo. Dying (A, upper panel) and EV^+^ (A, lower panel) cells are shown as % of PBMC. **(B)** Results from A) are plotted against groups HD (n = 30), RD (n = 12), mild (n = 15), moderate (n = 19) and severe disease (n = 15) for dying cells (upper panel) and EV^+^ cells (lower panel). **(C)** Plotting of the data from (B) against WHO ordinal scale. The plot shows the Spearman correlation test and linear regression line with 95% confidence interval shading (n = 38-79). **(D)** Summary of correlations of selected “nearest” (n = 23-49) laboratory and clinical parameters with peak WHO ordinal scale or and selected **(E)** our measurements of PS^+^, PS^+^EV^+^ and dying cells (bold) (n = 23-49). Significance was determined by Mann-Whitney test: *P < 0.05, **P < 0.01, ***P < 0.001, and ****P < 0.0001. Asterisks in brackets show statistically significant differences as compared to HD.

We confirmed these correlations in our patient cohort (Fig. Suppl. 2A). The frequencies of PS^+^EV^+^ and PS^+^ PBMC ranked in the top groups of measurements among inflammatory and coagulation parameters such as IL-6, PCT, CRP, PTT, D-Dimer, and others (Fig. Suppl. 2A). The strongest negative correlations existed with low lymphocyte counts (Fig. Suppl. 2A). For better comparability, we focused next only on those values determined from the same blood draw or close to our PS-measurements (‘nearest values’ in Fig. Suppl. 2A, shown in Fig. 2D). Here, both PS^+^ EV^+^ PBMC and PS^+^ PBMC correlated better than all other blood parameters with peak WHO ordinal scale (Fig. 2D). Moreover, PS^+^ EV^+^ PBMC frequencies correlated strongly withparameters of coagulation (fibrinogen, PTT), inflammation (IL-6, CRP), and lymphopenia (lymphocyte counts) (Fig. 2E, Fig. Suppl. 2B). Therefore, our new type of PS analysis allows the detection of subcellular particles associated with PBMC of COVID-19 patients. The percentage of PBMC bound to these PS^+^ particles correlated with the maximal (peak) WHO score of COVID-19 patients and allowed to classify patients with higher significance than previously established medical laboratory parameters^39^.

### PS^+^ EVs bind to several PBMC populations

Next, we investigated whether PS^+^ EVs would selectively associate with specific PBMC subpopulations. We examined CD4^+^ and CD8^+^ T cells (Fig 3A), CD19^+^ B cells (Fig. 3B), and HLA-DR^+^CD19^-^CD3^-^ PBMC (containing mainly monocytes and dendritic cells as central blood populations, Fig. 3B). In general, CD8^+^ T cells were more strongly associated with PS^+^ EVs than CD4^+^ T cells (Fig. 3A). However, both T cell subsets showed a similar tendency of increased PS^+^ EV binding in patients with a higher WHO score (Fig. 3A). B cells and blood monocyte-containing populations of patients in the different WHO groups were also associated relatively strongly with PS^+^ EVs (Fig. 3B). However, there was no actual grading with the severity of the disease. In summary, most blood cells showed a strong association with PS^+^ EVs in the patients. The frequency of PS^+^ EV^+^ CD8^+^ T cells best reflected the severity of COVID-19 disease.

**Fig. 3:**
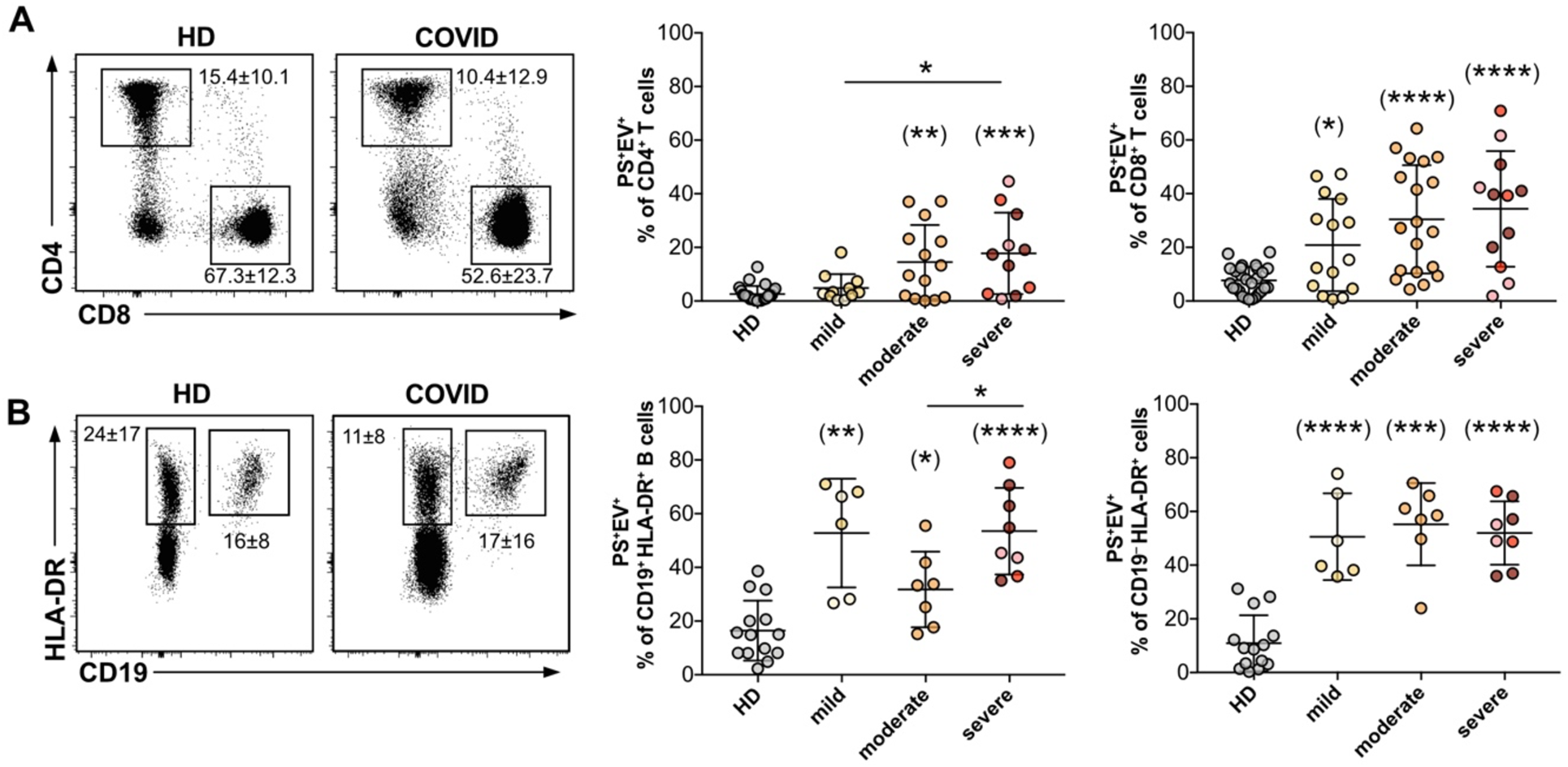
Identification of EV^+^ cells in PBMC from COVID-patients and healthy donors. PBMC were analyzed by IFC (gating strategy shown in Figure Suppl. 1A, B). PS^+^ CD4^+^ and CD8^+^ T cells **(A)** and CD19^+^ HLA-DR^+^ B cells and CD19-HLA-DR^+^ cells (containing mainly dendritic cells, monocytes) **(B)** were classified as PS^+^EV^+^ using the CAE and their total frequencies were plotted against HD (B, n = 14; A, CD4, n = 24; A, CD8, n = 27), mild (B, n = 6; A, CD4, n = 11; A, CD8, n = 15), moderate (B, n = 7; A, CD4, n = 14; A, CD8, n = 19), severe (B, n = 8; A, CD4, n = 11; A, CD8, n = 12) disease groups. Numbers next to the gates show the mean percentage ± SD of all cells depicted inside the dot plot that lie within the respective gate, while the graphs show the average frequency ± SD of EV^+^ cells within the analyzed subpopulation. Significance was determined by Mann-Whitney test: *P < 0.05, **P < 0.01, ***P < 0.001, and ****P < 0.0001. Asterisks in brackets show statistically significant differences as compared to HD.

### PS^+^ EVs associated with PBMC from COVID-19 patients are PS^+^CD41^+^ platelet-derived microparticles

EVs classify according to their size and origin into exosomes (up to 150nm), microvesicles or microparticles (MPs; 100-1000nm), and apoptotic EVs or apoptotic bodies (100-5000nm) ^40^. To better characterize the EVs associated with lymphocytes in COVID-19 patients, we isolated PS^+^ lymphocytes (B and T cells, PS^+^CD19^+^CD3ε^+^) from COVID-19 patients by FACS. We used recombinantly expressed, biotinylated MFG-E8-derived PS-binding domain mC1, which was multimerized by streptavidin (SA) (mC1-multimer/SA-APC) for PS-detection by flow cytometry (Fig. Suppl. 1C) and mC1-multimer/SA-gold for PS-detection by subsequent transmission electron microscopy (TEM). TEM pictures show cell-associated particles of a subcellular size bound to lymphocytes, which morphologically resemble T cells (Fig. 4 A-F).

**Fig. 4:**
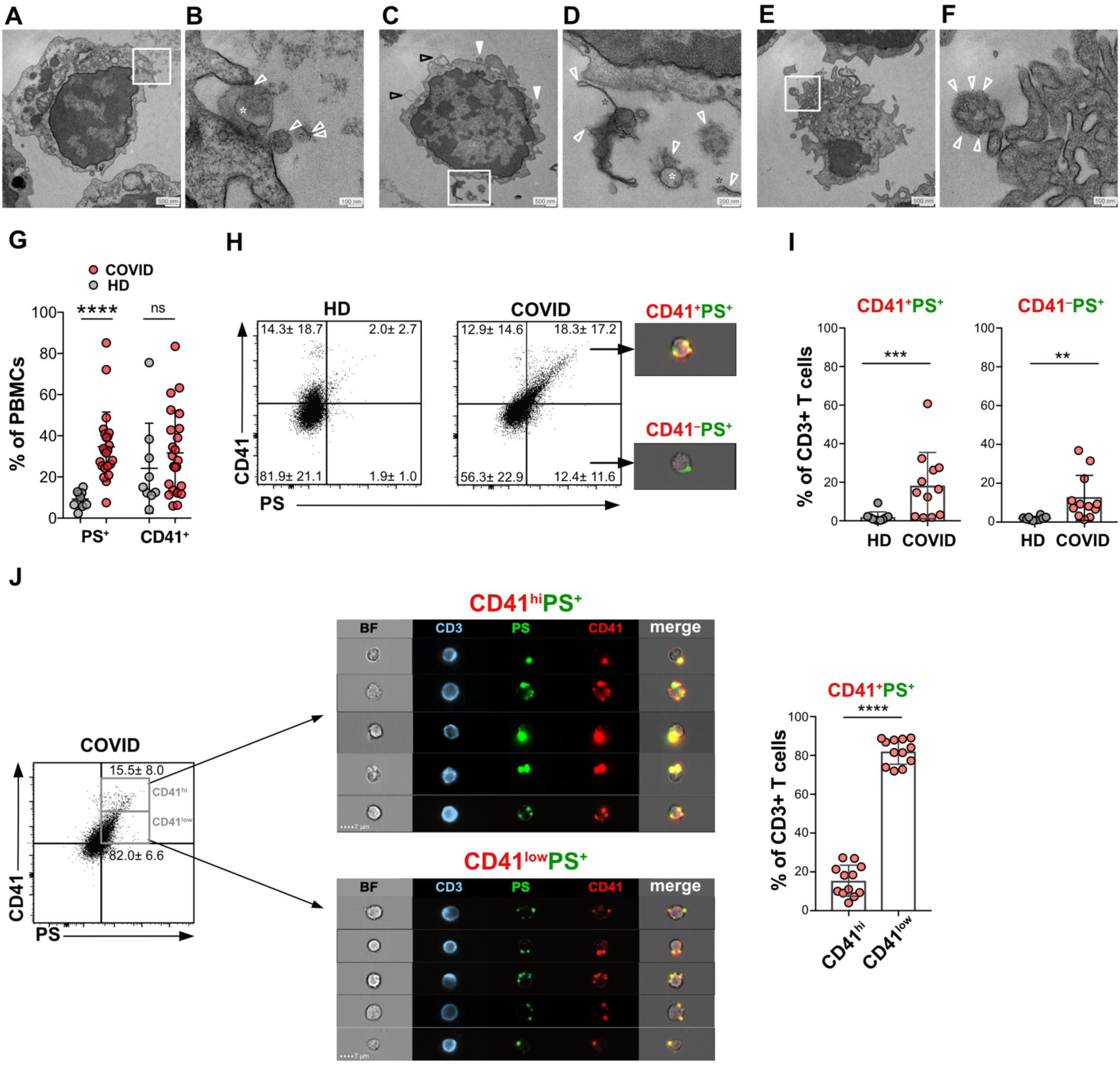
Structural and phenotypic characterization of EVs associated with lymphocytes from COVID patients. PS^+^ CD3ε^+^ T and CD19^+^ B lymphocytes were isolated from PBMC of COVID patients by FACS-sorting (Fig. Suppl. 1C) and subsequently labeled with mC1-multimer/streptavidin-gold for TEM analysis of PS and analyzed structurally by TEM **(A-F). (B, D, F)** represent the magnified sections (white frame) of **(A, C, E)**, respectively. Open white arrowheads (B, C, F) mark PS labeling by mC1-multimer/streptavidin-gold; arrowheads in C point to PS^−^ EVs; white star (B, D) or black star (D) marks organelle-like structures within EVs, or PS^+^ tubular elongated structures (D), respectively. Analysis of platelet marker CD41 on T cells. **(G)** PBMC were analyzed as shown in Fig. Suppl. 3 (HD, n =11; COVID, n =23) for PS and CD41. **(H)** CD3ε^+^ T lymphocytes (gated as in Fig. Suppl. 1D) were stained for CD41 and PS and analyzed by IFC (HD, n =10; COVID, n =12). Images represent cells of the respective quadrants. Numbers are the percentage of cells with the respective quadrants. **(I)** Percentage of T cells in the quadrants of CD41^+^PS^+^ and CD41^-^PS^+^ cells. **(J)** Dot plots show the gating of PS^+^ CD41^hi^ and PS^+^CD41^low^ CD3ε^+^ T lymphocytes. IFC images show representative cells of the CD41^hi^ and CD41^low^ gates. The bar graph shows the percentage of PS^+^CD41^hi^ and PS^+^CD41^low^ T cells. Statistical significance was determined by paired Wilcoxon test and is indicated by asterisks (ns P > 0.5; *P ≤ 0.05; **P ≤ 0.01; ***P ≤ 0.001; two-tailed unpaired t-test).

Many (Fig. 4B, D, F, open white arrows), but not all EVs (Fig. 4C, arrows), were PS^+^, and most EVs were >100 nm in size (Fig. 4B, D, F). The particles were mainly round and had different shapes and contained other smaller components, cell contents, or organelles (Fig. 4B, D, white star). However, we also found PS^+^ tubular elongated structures (Fig. 4D, black star). Relatively large particles with highly diverse shapes, including tubular shapes, and content are known features of PMPs, 50% of which are PS positive ^41-43^. Hyperactivated platelets in COVID-19 patients ^44^ are sources for PMPs and contribute to the hypercoagulability state of the disease ^45-47^. Activated platelets can release two types of EVs, (i) smaller exosomes (40-100 nm in diameter) derived by exocytosis from α-granules and the multivesicular body, and (ii) PS^+^ MP (100-1000 nm in size), formed by budding of the plasma membrane ^48,49^. These similarities let us hypothesize that PMPs attach to PBMC of COVID-19 patients.

To test this hypothesis, we stained PBMC from COVID-19 patients for CD41, a platelet marker, part of a fibrinogen-receptor, and present on platelet-derived PMPs ^48^ (Fig. Suppl. 3, Fig. 4G). Analysis of flow cytometry data of PBMC confirmed our assumption and showed that many PBMC were positive for the platelet marker CD41 (Fig. 4G). However, CD41 could not distinguish PBMC from healthy donors and COVID because CD41^+^ PBMC also existed in healthy donors (Fig. 4G, CD41^+^). In contrast, PS-positivity alone distinguished PBMC from healthy donors and COVID-19 patients, as PS^+^ PBMC were highly significantly enriched in patients (Fig. 4G, PS^+^).

To analyze PS and CD41 and their possible colocalization, we performed image analysis of PBMC (Fig. Suppl. 3A, B) and T lymphocytes (Fig. 4H-J). In COVID-19 patients, we observed a substantial increase of PS and CD41 double-positive PBMCs and T cells from 10 to 40% and from 2 to 23%, respectively. The increase in CD41^-^PS^+^ cells was not as pronounced but still significant (Fig. 4H-I, Fig. Suppl. 3A, B). This finding shows that most PS^+^ cells acquire PS through the binding of CD41^+^ platelets or their PMPs. To investigate whether the cells preferentially bind whole platelets or smaller PMPs, we quantified CD41^hi^ versus CD41^low^ cells. The CD41^hi^ gate contained mainly cells with large CD41^+^ particles attached, which were also visible in the brightfield (BF) channel – presumably whole platelets. In contrast, the cells in the CD41^low^ gate were associated with small, more dimly stained spots that were too small to be visible in the BF channel - presumably PMPs (Fig. 4J). While T cells very clearly had a strong preference for PMP binding (80% of PS^+^ T cells were CD41^low^, Fig. 4J), PBMCs did not show such a clear preference (approx. 50% were CD41^low^, Suppl. Fig. 3B). Our data show that PBMCs from COVID-19 patients associate to a high degree with PS^+^ platelets and PMPs, while T cells rarely bind whole platelets, but rather PMPs.

### PMPs carry markers of activated platelets

SARS-CoV-2-infection causes platelet activation and alters their functions ^50,51^. The results of our study suggested that the PMPs associated with lymphocytes in COVID-19 patients originate from activated CD41^+^ platelets. In addition to PS, markers such as CD62P (P-selectin) and the tetraspanin CD63 are present on PMPs derived from activated platelets ^52^. Our patient cohort showed increased percentages of PS^+^ and CD62P^+^ platelets, both markers for platelet activation (Fig. Suppl. 4A-C). When associated with CD8^+^ T cells, platelet-derived molecules PD-L1 (CD274) ^53^ and CD86 ^54^ on PMPs could be potentially negative or positive costimulatory triggers, respectively. Platelets were strongly positive for CD274 (Fig. Suppl. 4B, C), but this was not different between healthy donors and patients (Fig. Suppl. 4B). The same was true for CD86, which we found in similar amounts and generally only very few platelets that were positive for this molecule in both groups (Fig. Suppl. 4B, C). We evaluated next if PS^+^EV^+^ CD8^+^ T cells were positive for these platelet markers. PS^+^EV^+^ CD8^+^ T cells showed significantly increased mean fluorescence intensities (MFI) for platelet markers CD41, CD63, CD62P, and CD274 compared to their PS^-^ counterparts (Fig. 5A, B). The frequency of T cells, which were positive for these markers, was also increased in PS^+^EV^+^ T cells (CD41, CD63, CD62P; Fig. 5B). This finding indicated that PMPs carried the surface molecules of their activated ‘parent’ platelets to the surface of activated CD8^+^ T cells in COVID-19 patients. When we analyzed the images of CD41, CD63, CD274, and CD62P stained T cells, we also observed an EV-like staining pattern of these markers, similar to the PS staining (Fig. 5C). This finding further confirms that T cells can acquire these markers through vesicles. To confirm that these markers originate from activated platelets or their PMPs, we quantified PMP-marker colocalization with PS. We used the bright detail similarity (BDS) feature to quantify the pixel overlap of PS^+^ and PMP-marker^+^ spots identified using a peak mask in the IDEAS software. BDS values >1 indicate substantial colocalization between PS and the other markers. This analysis showed that the great majority of all PS^+^EV^+^ CD8^+^ T cells (64% - 82%) showed strong colocalization of PS with CD41, CD63, CD62P, or CD274 with average BDS scores ranging from 1.3 to 1.8 (Suppl Fig. 5). These results indicate that most of the PMPs detected on CD8^+^ T cells originate from activated platelets and carried platelet markers.

**Fig. 5.**
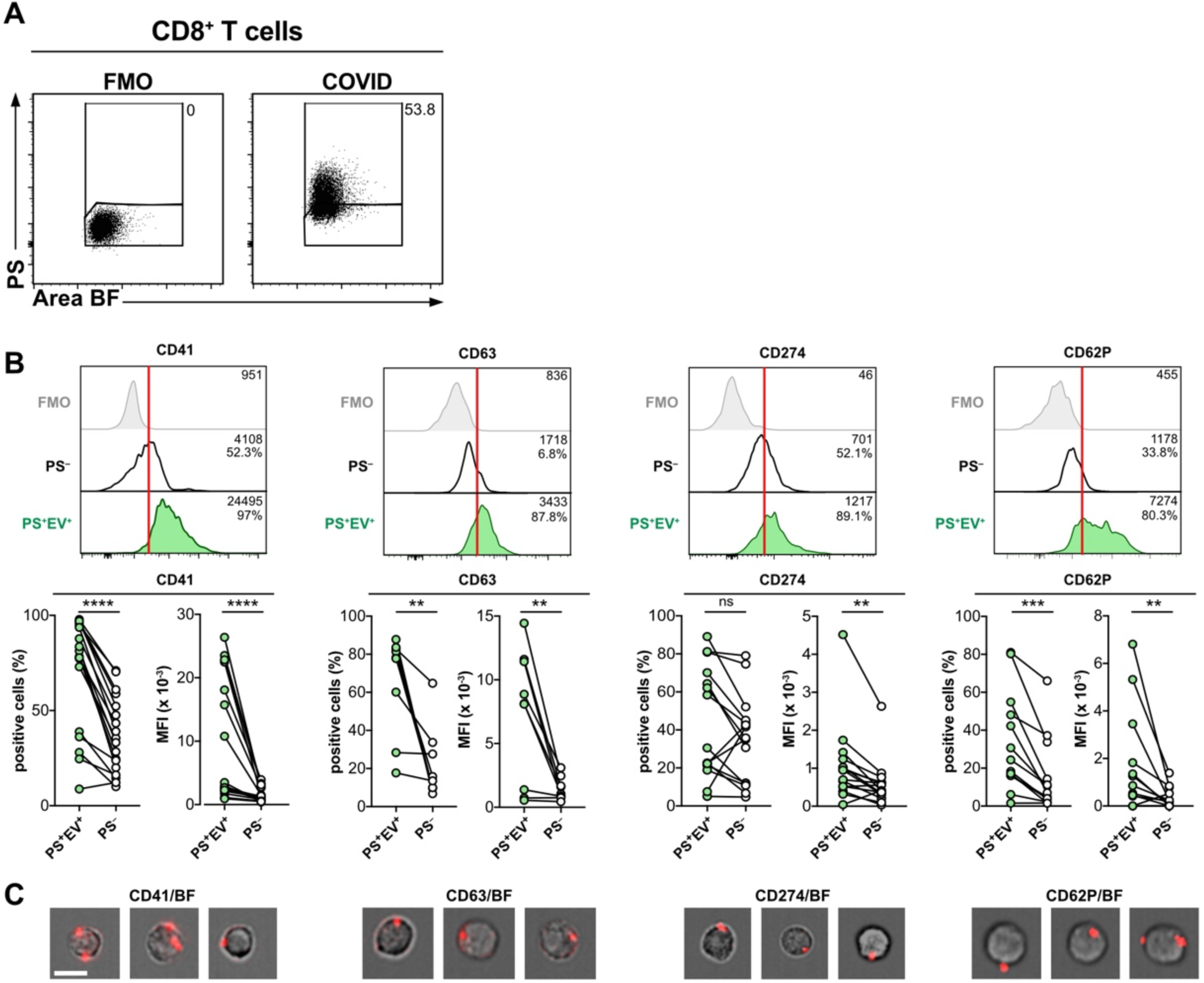
EVs bound to CD8^+^ T cells originate from platelets. **(A)** Analysis of CD8^+^ T cells of COVID patients (CD41, n = 18; CD63, n = 9; CD274, n = 15, CD62P, n = 13) were gated as shown in Fig. Suppl 1E and then further separated into PS^+^EV^+^ and PS^-^ T cells. Numbers represent the percentage of cells within the gates. **(B)** Then cells were analyzed separately for expression of platelet-markers CD41, CD63, CD62P, and CD274. Median fluorescence intensities (MFI) ± SD of these proteins and % of positive cells are indicated in the histograms. Summaries of all results are shown below histograms. **(C)** Representative images of CD41^+^, CD63^+^, CD62P^+^, and CD274^+^ CD8^+^ T cells show the EV-like staining pattern of these markers. Statistical significance was determined by paired Wilcoxon test and is indicated by asterisks (ns P > 0.5; *P ≤ 0.05; **P ≤ 0.01; ***P ≤ 0.001).

### PS^+^CD41^+^ PMPs are preferentially bound to proliferating CD8^+^ T cells

Next, we wanted to assess whether there are functional differences between PMP^+^ and PMP^-^ CD8^+^ T cells. For this, we performed RNAseq analysis of FACS-sorted PS^+^ and PS^-^ non-naïve CD8^+^ T cells from peripheral blood of 4 patients (Fig. 6A, Suppl. Fig. 1G). Despite heterogeneity between patients, we could identify gene sets that showed clear enrichment in PS^+^ (Fig. 6B) and PS^-^ T cells (Fig. 6C).

**Fig. 6:**
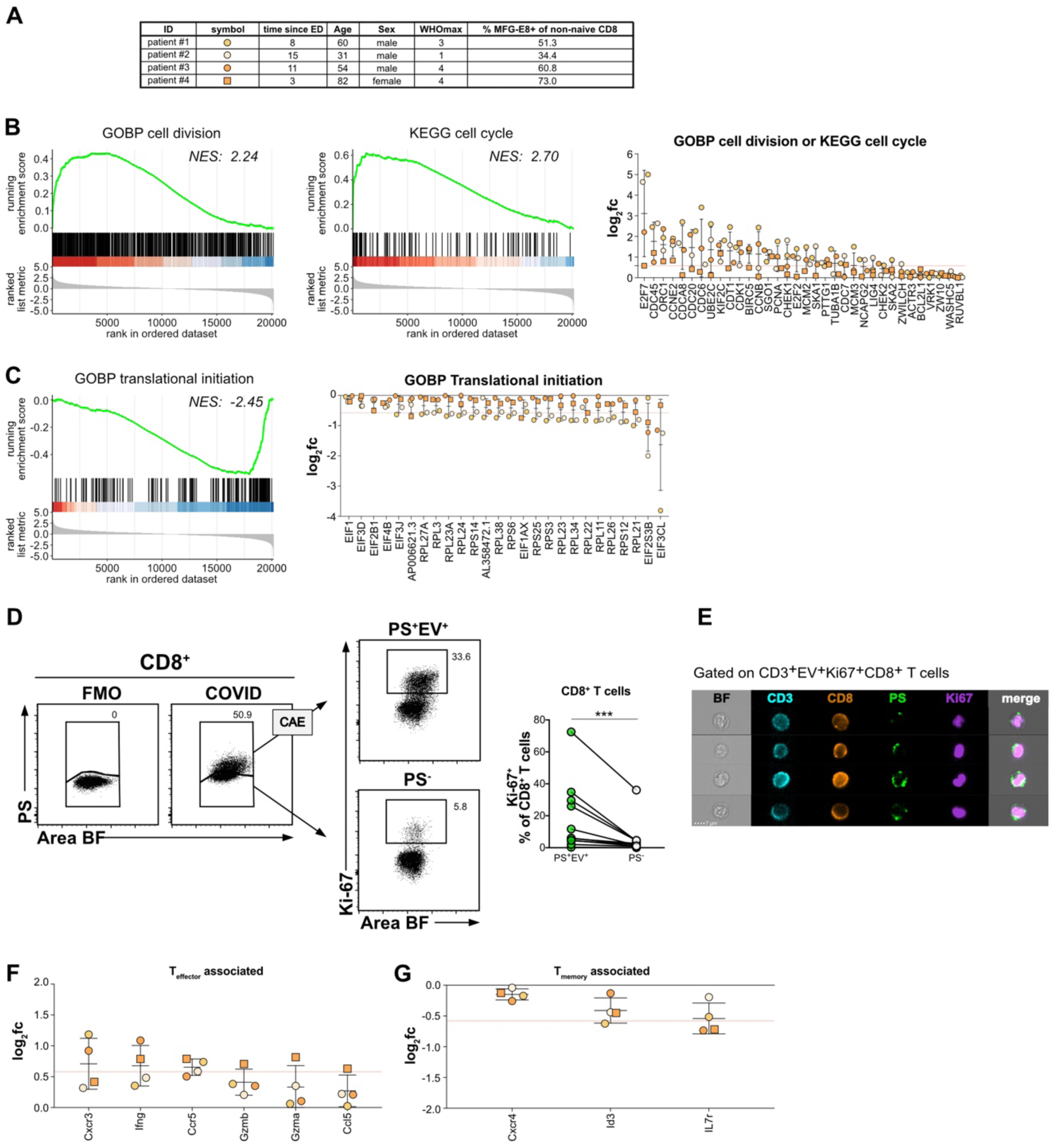
PS^+^EV associate with proliferating CD8 T cells. Non-naive PS^+^ and PS^-^ CD8^+^ T cells (gating strategy see Suppl. Fig. 1G) from 4 patients were sorted and subjected to RNAseq analysis. **(A)** The table displays patient parameters. **(B)** GSEA enrichment analysis for the gene sets GOBP cell division and KEGG cell cycle are shown and indicate enrichment in PS^+^ CD8 T cells. Dot plots depict log2 fold change values of genes in either of these two gene sets and upregulated in all four patients. The red dotted horizontal line indicates a fold change of 1.5 (log2fc 0.58). **(C)** GSEA enrichment analysis for the gene sets GOBP translational initiation and indicates enrichment in PS^-^ CD8^+^ T cells. Dot plots depict log2 fold change values of genes in this gene set and downregulated in all four patients. The red dotted horizontal line indicates a fold change of -1.5 (log2fc -0.58). **(D)** PBMC from COVID patients were examined for proliferation with Ki-67. CD8^+^ T cells (n = 11) were analyzed as shown in Figure Suppl. 1F. PS^+^CD3^+^CD8^+^ T cells were stained intranuclear for Ki-67. PS^+^ and PS^−^ fractions were classified by IFC. CAE-analysis identified PS^+^EV^+^ live cells. PS^+^EV^+^ and PS^-^ CD8^+^ T cells were then analyzed for Ki-67, and data from all patients were plotted on the graph. The numbers in the dot plots represent the fraction of cells in the corresponding gate. Statistical significance was determined by paired Wilcoxon test and is indicated by asterisks (ns P > 0.5; *P ≤ 0.05; **P ≤ 0.01; ***P ≤ 0.001). **(E)** shows an image selection of CD8 T cells with the markers used. **(F)**-**(G)** Log2 fold change values of genes associated with T effector (F) and T memory (G) cells that were up- or downregulated, respectively, in all four patients, are shown. Red dotted horizontal line indicates a fold change of 1.5 (log2fc 0.58) or -1.5 (log2fc -0.58).

PS^+^ CD8^+^ T cells had enriched genes controlling cell cycle and division, with normalized enrichment scores (NES) of 2.24 and 2.70 for the gene ontology biological process (GOBP) ‘cell division’ and KEGG’ cell cycle’ gene sets, respectively. Among the genes upregulated in PS^+^ CD8^+^ T cells of all patients were several cell division control (CDC) genes, such as CDC45, CDCA8, CDC20, CDC6, or transcription factors E2F7 and E2F2, which are involved in cell cycle regulation ^55^. Also, Birc5 (survivin) which plays a crucial role in costimulation-driven clonal expansion of T cells ^56^, was upregulated in all patients.

Cells in the G2/M phase exhibit a general inhibition of translation ^57^. Hence, the reduced expression of genes initiating translation in PS^+^ CD8^+^ T cells is in line with the observed upregulation of proliferation-associated genes (Fig. 6C).

Although PS^-^ and PS^+^ CD8^+^ T cells had few apparent differences in gene expression due to the high degree of variability between individual patients, it is striking that binding of PS^+^ PMPs was associated with increased proliferation. To confirm the RNAseq results, we stained CD8^+^ T cells with the proliferation marker Ki-67, which labels dividing and recently dividing cells in G1, S, G2, and M phase, but is absent in resting cells ^58^ and compared the frequency of PS^-^ and PS^+^EV^+^ Ki-67^+^ cells. PS^+^EV^+^ CD8^+^ T cells (Fig. 6D, E) and PS^+^EV^+^ CD4^+^ T cells (Fig. Suppl. 6) contained significantly more Ki-67^+^ proliferating cells than their PS^-^ T cell counterparts from the same patient. In both cases, Ki-67 staining localized to the nucleus and PS^+^ PMPs to the periphery of the same cells (Fig. 6E, Fig. Suppl. 6B). Our results indicate that PS^+^ PMPs preferentially bind to proliferating T cells and may affect T cells in this cycling stage.

We were also interested to find out, whether CD8^+^ T cells binding PS^+^ PMPs exhibit an effector or memory phenotype. Previous reports showed a high prevalence of SARS-CoV-2-specific T cells mainly among T cells with phenotypes of effector memory (TEM) and terminal effector memory cells reexpressing CD45RA (TEMRA) cells ^59^. We did not observe a clear enrichment of effector or memory genes ^60^. However, we found an upregulation of effector-associated genes, such as CXC Motif Chemokine Receptor 3 (Cxcr3), Interferon γ (Ifng), Granzyme A and B (Gzma and Gzmab), and CC-chemokine ligand 5 (Ccl5) in PS^+^ CD8^+^ T cells of all four patients. In contrast, memory-related genes, such as C-X-C Motif Chemokine Receptor 4 (Cxcr4), DNA-binding protein inhibitor ID-3 (Id3), and interleukin-7 receptor (Il7r), showed a subtle downregulation in all patients. These findings indicate that PMP-associated CD8^+^ T cells are proliferating effector cells rather than memory cells. This we also confirmed with flow cytometry using CCR7 and CD45RA as markers ^61^. We found PS^+^PMPs preferentially associated with CCR7^-^CD45RA^-^ CD8^+^ TEM cells and terminally differentiated CCR7^-^CD45RA^+^ CD8+ TEMRA.

## DISCUSSION

COVID-19 patients have hyperactivated platelets ^50^, elevated levels of circulating PMPs ^62^, and an increased risk of thromboembolic complications contributing to disease severity and mortality.

Our results contribute to the complex clinical picture of thromboinflammation (reviewed in ^4,36^). One of our most surprising findings was the high association of PBMC with PS^+^ PMPs and platelets over the disease course, shown with a novel PS-detecting method. The degree of this association correlated more strongly with disease severity than established laboratory indices such as lymphopenia, IL-6, D-Dimer, fibrinogen, and others measured simultaneously. Antiphospholipid autoantibodies (aPL antibodies), including autoantibodies to the PS/prothrombin complex (antiPS/PT), have been found in COVID-19 patients^6^. Hence, it is possible that the association of PS^+^ platelets and PMPs with activated lymphocytes, together with a highly inflammatory environment, may drive the generation of aPL antibodies, further increasing the risk of life-threatening thrombotic events.

Patients with COVID-19 are mostly lymphopenic^18,19^, but lymphopenia preferentially affects CD8^+^ T cells ^23,63^. The measurement of dying or apoptotic cells *in vivo* or fresh *ex vivo* samples is technically challenging. Phagocytes very efficiently remove dead cells, and sample preparation itself can contribute to cell death. Using novel PS-specific reagents based on lactadherin, we found few but statistically significantly increased amounts of dying cells in the blood of COVID-19 patients with mild (WHO 1-3) and moderate disease (WHO 4). However, it is currently unclear, why PBMC death occurs during these relatively mild stages of the disease.

By interacting with the complement cascade activated in COVID-19 patients ^64^, PS^+^ PMPs could contribute to lymphopenia. Correlation analyses of all blood values with the amounts of PS^+^PMP^+^ PBMCs showed the strongest negative correlation with decreased lymphocyte counts in patients’ blood, i.e., with established lymphopenia. PS and PS^+^ PMP can activate complement pathways ^7-9^, thrombin formation, coagulation, and inflammation ^12-14^. PS^+^PMP^+^ T cells could trigger for example the complement cascade on their surface, which might cause cell damage, death, and T cell removal by phagocytes.

Moreover, the presence of an activated complement system in COVID-patients ^64^ could increase the release of PMPs from platelets ^65^ to self-reinforce this spiral. PS^+^ PMPs might facilitate the assembly and dissemination of procoagulant enzyme complexes ^65^. Attracting these reactions to the surface of lymphocytes might cause further functional inhibition or cell death.

“Long Covid” is a phenomenon that occurs in around 10% of COVID-19 patients and seems to be associated with persistent tissue damage in severe cases. However, also patients with mild COVID-19 disease scores might be affected ^66^. We identified PS^+^PMP^+^ PBMCs of patients during many weeks post initial diagnosis with only minimal signs of returning to normal levels. Therefore, prolonged adverse effects of PS^+^ PMPs on the immune system could contribute to “long COVID".

Especially CD8^+^ TEM and TEMRA cells, among which most SARS-CoV2-specific clones were identified ^59^ showed enhanced PMP-binding in COVID-19 patients. COVID-19 T cell responses begin too late ^30^, and T cells may be exhausted ^67^. The association of PS^+^ PMPs with T cells could contribute to these deficiencies and potentially impact the immune and inflammatory antiviral responses. Activated platelets may associate with T cells in the blood of HIV-infected individuals ^68^ and link the coagulation and inflammatory cascade with T cells. *In vitro*, platelets can inhibit proliferation, cytokine production, and PD1 expression of T cells ^69^. Since PMPs derive from platelets, they may also have similar functions as their ‘parent’-platelets. We found that between 10-80% of CD8^+^ T cells were associated with PD-L1(CD274)^+^ PMP. The fact that we could detect this immunoregulatory molecule on a high frequency of PS^+^ and PS^-^ PMPs raises the possibility that PMPs can suppress CD8^+^ T cells via PD-L1/PD1 interaction. Previous studies suggested that SARS-CoV-2-specific CD8^+^ T cells can have an exhausted phenotype due to their expression of inhibitory receptors such as PD1 ^67,70-72^. Analogous to PD-L1 on tumor exosomes, which suppress tumor-specific CD8^+^ T cells^73^, PD-L1 PMPs could favor T cell suppression in COVID-19. However, recent data have suggested that PD1^+^ SARS-CoV-2-specific CD8^+^ T cells are functional ^74^. Furthermore, our RNAseq results showing subtly increased expression levels of the effector molecules IFNγ, granzyme A, and B and increased proliferation also argue against an exhausted phenotype of PS^+^ CD8^+^ T cells and would instead indicate enhanced effector function or terminal differentiation.

Two recent studies showed that in COVID-19 patients, hyperactivated platelets could form aggregates with leukocytes and macrophages ^51,75^. As these previous studies relied on conventional flow cytometry, but not IFC, they could not differentiate between CD41^+^ platelets and CD41^+^ PMPs. Also, they did not analyze PS^+^ PMPs on the surface of live lymphocytes. Since PMPs constitute the lion’s share of CD41^+^PS^+^ particles on T cells of COVID-19 patients and PS may be responsible for the coagulation and inflammatory effects mentioned above, our findings are an unexpected result with high clinical relevance.

PS is a novel marker to classify COVID-19 patients according to disease severity. Moreover, longitudinal studies could test PS as a predictor of disease development. Due to its ease of use by flow cytometry and the high number of positive PS^+^ PBMC during COVID-19, PS detection could be a valuable analytical tool also in other infectious diseases and sepsis.

## ACKNOWLEDGMENTS

We would like to thank all of the COVID-19 Registry of the LMU Munich (CORKUM) investigators and staff and the patients and their families for participating in the CORKUM registry and all health care workers for their outstanding service. We acknowledge the Core Facility Flow Cytometry at the Biomedical Center, LMU Munich, for providing the ImageStreamX MKII imaging flow cytometer. T.B. and A.K. are supported by the Deutsche Forschungsgemeinschaft under CRC 1054 (TP B03, grant no. 210592381; and TP A06, grant no. 210592381). A.K. received funding from Deutsche Forschungsgemeinschaft under CRC/TR237-B14 (369799452) and KR2199/10-1 (391217598), and received funding from the Bavarian State Ministry of Science and the Arts.

## AUTHOR CONTRIBUTIONS

Conceptualization, J.K. and T.B.; Software, J.K.; Formal analysis, T.S., L.Ra., K.S., M.S., M.Sch.; Investigation, L.Ra., R.G., L.Ri., M.Sch. and J.K.; Resources, J.H., C.S., M.M., E.W., J.C.H, M.B.B.; Writing – Original draft, T.B., J.K., and L.Ra; Writing – Review & Editing, A.K. M.S., M.Sch., M.M., R.G.; Supervision, T.B., A.K., J.K.; Funding Acquisition, T.B., A.K., M.S.

## MATERIALS AND METHODS

### Study design and recruitment

Recruited patients (n=54) with PCR confirmed SARS-CoV-2 infection were part of the COVID-19 Registry of the LMU University Hospital Munich (CORKUM, WHO trial id DRKS00021225). The Ethics Committee approved the study of the LMU Munich (No: 20-308; 18-415), and patients included (≥ 18 years, mean age 63, Suppl. Table 1) consented to serial blood sampling. Additional approval was obtained for the analysis shown here (No. 20-308) and for the use of blood samples from healthy donors (No. 18-415). For this study, patient data were anonymized for analysis, and blood samples were collected between April 2020 and February 2021. Of the 54 patients analyzed, 52 patients were hospitalized, and two patients were diagnosed in the hospital and discharged. From 15 patients, several timepoints could be obtained, which were taken between 2 and 76 after a positive SARS-CoV-2 PCR test. Clinical and laboratory data were collected and documented by the CORKUM study group and are summarized in Supplemental Table 1. We used the World Health Organization’s (WHO) eight-point ordinal scale for COVID-19 endpoints ^37^ to grade disease severity. WHO scores were combined into ‘mild‘ (WHO 1-3), ‘moderate’ (WHO 4), and ‘severe’ (WHO 5-8). Recovered donors (RD, n=12, mean age 40, Suppl. Table 2) were adults with a prior SARS-CoV-2 infection (≥ 69 days post positive PCR test), who were either diagnosed in the ambulance or released from the hospital with WHO score 1-2.

Healthy donors (HDs, n=35, mean age 39, Suppl. Table 3) tested negative for SARS-CoV-2 were used as control cohorts. PBMCs were obtained from leucocyte reduction chambers after thrombocyte donations (n=10) or freshly prepared from whole blood (n=25) of healthy hospital and laboratory workers.

### Sample processing and cell isolation

Peripheral blood was collected into lithium heparin tubes (Sarstedt) and processed within 6 hours after venipuncture. Unfixed samples were handled under Biosafety level 2. PBMCs were isolated by Biocoll density gradient (Merck) centrifugation and then directly stained for imaging flow cytometry.

### Antibody staining and flow cytometry

Antibodies and staining reagents are listed in Suppl. Table 4. As PS-staining reagents we used mC1-biotin multimerized with streptavidin-AF647 for (image) flow cytometry or streptavidin-gold for TEM. mC1-multimers are commercially available through BioLegend. In flow cytometry we also used the previously published recombinant MFG-E8-eGFP ^33,34^. All PS-staining reagents are Ca^2+^ independent, highly stable and PS-specific. Freshly isolated PBMCs were incubated with Fc block before live/dead and cell surface antibody staining. PBMCs surface staining was performed in staining buffer (PBS containing 2% of fetal calf serum) for 25 min on ice. Then cells were washed and fixed with 4 % PFA for 30 min at room temperature (RT), washed again, filtered, and analyzed on an ImageStreamx MKII imaging flow cytometer (Luminex). Samples were acquired with low flow rate/high sensitivity and 60x magnification. Images were acquired using bright field illumination and the excitation lasers 405, 488, 561, and 642 nm.

For intranuclear Ki-67 staining, cells were fixed and permeabilized using the Transcription Factor Staining Buffer Set (ThermoFisher, cat. #00-5523-00) according to the manufacturers’ instructions. Intranuclear staining was performed in permeabilization buffer for 30 min at RT.

### Cell sorting for gene expression analysis

PBMCs from four patients (see Fig. 6 A) were stained with anti-CD3-BV421, anti-CD8-BV785, anti-CCR7-APC, anti-CD45RA-PerCPCy5.5, anti-CD19/CD16/CD56/CD14-APCFire750 and PS-reagent (see Suppl. Table. 4) in staining buffer (25 min, 4C°) after cells were incubated with Fixable Viability dye eFluor780 in PBS (10 min, 4°C). PS^+^ and PS^-^ single, live, non-naïve (CCR7^+^CD45RA^-^, CCR7^-^CD45RA^-^ and CCR7^-^CD45RA^+^) CD3^+^CD8^+^ T cells were directly sorted into TRIzol reagent (Thermo Fisher) on a FACSAriaFusion (BD) using a 100 μm nozzle. Samples were stored at – 20 °C until analysis. RNA isolation and sequencing were performed by Vertis Biotechnologie AG, Freising. Total RNA was isolated and purified using Monarch columns (NEB). Poly(A)+ RNA was isolated from the total RNA sample. First-strand cDNA synthesis was primed with a N6 randomized primer. After fragmentation, the Illumina TruSeq sequencing adapters were ligated in a strand specific manner to the 5’ and 3’ ends of the cDNA fragments. In this way, a strand specific PCR amplification of the cDNA was achieved using a proof-reading enzyme. For Illumina NextSeq sequencing, the samples were pooled in approximately equimolar amounts. The cDNA pool in the size range of 250-700 bp was eluted from a preparative agarose gel. The primers used for PCR amplification were designed for TruSeq sequencing according to the instructions of Illumina. The cDNA size range was 250-700 bp. The cDNA pool was sequenced on an Illumina NextSeq 500 system using 1×75 bp read length.

### RNAseq analysis

Sequencing reads were aligned to the human reference genome (version GRCH38.100) with STAR (version 2.7.3). Expression values (TPM) were calculated with RSEM (version 1.3.3). Post-processing was performed in R/bioconductor (version 4.0.3) using default parameters if not indicated otherwise. Differential gene expression analysis was performed with ‘DEseq2’ (version 1.28.1) using a model including patient ID as random factor. An adjusted p value (FDR) of less than 0.1 was used to classify significantly changed expression. Gene set enrichment analyses were conducted with ‘clusterProfiler’ (version 3.18.1) on the statistic reported by DEseq2. Data are available at GEO Submission (GSE174786).

### Cell sorting for transmission electron microscopy (TEM)

Isolated PBMCs were stained with anti-CD19- and anti-CD3-PE, PS-staining reagent mC1-multimer (SA-AF647) (commercially available at BioLegend), and mC1-biotin (commercially available at BioLegend) multimerized with SA-gold, (Aurion) (see Suppl. Table 4) in staining buffer (25 min, 4 °C) after incubation with Fixable Viability Dye eFluor780 in PBS (10min, 4°C). After washing, cells were prefixed with freshly prepared 4% EM-grade PFA (Science Services) for 30 min at RT before sorting. Fixed, single, PS^+^CD19^+^ and CD3^+^ cells were sorted on a FACSAriaIII (BDBiosciences) using a 130 μm nozzle into PBS containing 0.5 % BSA. After washing with PBS, cells were fixed with 2.5 % glutaraldehyde (EM-grade, Science Services) in 0.1 M cacodylate buffer (pH 7.4) for 15 min (Sigma Aldrich) and washed with 0.1 M sodium cacodylate buffer for 10 min at 400g before postfixation in reduced osmium (1% osmium tetroxide (Science Services), 0.8 % potassium ferricyanide (Sigma Aldrich) in 0.1 M sodium cacodylate buffer). The cell pellet was contrasted in 0.5 % uranyl acetate in water (Science Services and dehydrated in an ascending ethanol series. The pellet was embedded in epon resin and hardened for 48 hours at 60°C. Ultra-thin sections (50 nm) were cut and deposited onto formvar-coated grids (Plano) and again contrasted using 1 % uranyl acetate in water and Ultrostain (Leica). Images were acquired on a JEM 1400plus (JEOL).

### Imaging flow cytometry and data analysis

Data analysis was performed using the IDEAS software (Version 6.2, Luminex). Compensation matrices were generated using single stained samples and applied to the raw data, and data analysis files were created. Unfocused events were excluded from the analysis based on gradient max feature values. PS^+^ cells were gating using FMO controls. TIFF-images of PS^+^ cells from each sample were exported (16-bit, raw) and analyzed by the CAE algorithm as previously described ^33^. Two *.pop files containing the object numbers of apoptotic and EV^+^ cells were generated and re-imported into the IDEAS software. Then FCS files from all cells, only apoptotic or only EV^+^ cells, were exported and analyzed using FlowJo Version 10.7.1.

### Colocalization analysis

We created a spot mask for PS and the respective marker staining to determine the degree of colocalization between PS^+^ EVs and PMP-marker^+^ (CD41/CD63/CD274/CD62P) cells. PS^+^ spots were identified by the Dilate(Peak(M02, PS, Bright, 5),1) mask and the Dilate(Peak(M_marker, marker channel, Bright, 1),1) mask was used to identify marker^+^ spots. Both masks were combined, and the colocalization was assessed using the BDS R3 feature of the IDEAS software.

### Platelet staining

Platelets were stained in whole blood. 100 μl of blood were slowly added to 100 μl of antibody mix in staining buffer (PBS with 0.1% sodium azide and 2 % of FCS). The mixture was gently swirled and incubated for 30 min at room temperature in the dark. Then cells were fixed in 1% PFA in PBS with 0.1% sodium azide for 2 hours, centrifuged for 20 min at 5000g, resuspended in PBS with 0.1% sodium azide, and analyzed on the ImageStreamx MKII imaging flow cytometer.

### Statistical analysis

For statistical analysis, the PRISM software (GraphPad Software, La Jolla, CA, USA) was used. For direct comparison between two groups non-parametric, unpaired Mann-Whitney test was used. Statistical significance of paired data was determined by paired Wilcoxon test. P values of ≤0.05 are considered significant and donated with *, **P < 0.01, ***P < 0.001, and ****P < 0.0001. R version 4.0.3 was used for correlation analysis. Numeric values in the dataset were correlated (Spearman correlation) using the ggcorrmat function of the ggstatsplot package (v0.7.2) with Benjamini-Hochberg correction for multiple testing.

## REFERENCES

1 Guan, W. J. et al.. Clinical Characteristics of Coronavirus Disease 2019 in China. N Engl J Med 382, 1708–1720, doi:10.1056/NEJMoa2002032 (2020).

2 MacLaren, G., Fisher, D. & Brodie, D. Preparing for the Most Critically Ill Patients With COVID-19: The Potential Role of Extracorporeal Membrane Oxygenation. JAMA 323, 1245–1246, doi:10.1001/jama.2020.2342 (2020).

3 Wolfel, R. et al.. Virological assessment of hospitalized patients with COVID-2019. Nature 581, 465–469, doi:10.1038/s41586-020-2196-x (2020).

4 Gu, S. X. et al.. Thrombocytopathy and endotheliopathy: crucial contributors to COVID-19 thromboinflammation. Nat Rev Cardiol 18, 194–209, doi:10.1038/s41569-020-00469-1 (2021).

5 Corban, M. T. et al.. Antiphospholipid Syndrome: Role of Vascular Endothelial Cells and Implications for Risk Stratification and Targeted Therapeutics. J Am Coll Cardiol 69, 2317–2330, doi:10.1016/j.jacc.2017.02.058 (2017).

6 Zhang, Y. et al.. Coagulopathy and Antiphospholipid Antibodies in Patients with Covid-19. The New England journal of medicine 382, e38, doi:papers3://publication/doi/10.1056/NEJMc2007575 (2020).

7 Wang, R. H., Phillips, G., Jr., Medof, M. E. & Mold, C. Activation of the alternative complement pathway by exposure of phosphatidylethanolamine and phosphatidylserine on erythrocytes from sickle cell disease patients. J Clin Invest 92, 1326–1335, doi:10.1172/JCI116706 (1993).

8 Mevorach, D., Mascarenhas, J. O., Gershov, D. & Elkon, K. B. Complement-dependent clearance of apoptotic cells by human macrophages. J Exp Med 188, 2313–2320, doi:10.1084/jem.188.12.2313 (1998).

9 Tan, L. A., Yu, B., Sim, F. C., Kishore, U. & Sim, R. B. Complement activation by phospholipids: the interplay of factor H and C1q. Protein Cell 1, 1033–1049, doi:10.1007/s13238-010-0125-8 (2010).

10 Huong, T. M., Ishida, T., Harashima, H. & Kiwada, H. The complement system enhances the clearance of phosphatidylserine (PS)-liposomes in rat and guinea pig. International Journal of Pharmaceutics 215, 197–205, doi:10.1016/s0378-5173(00)00691-8 (2001).

11 Paidassi, H. et al.. C1q binds phosphatidylserine and likely acts as a multiligand-bridging molecule in apoptotic cell recognition. J Immunol 180, 2329-2338 (2008).

12 Owens, A. P., 3rd & Mackman, N. Microparticles in hemostasis and thrombosis. Circ Res 108, 1284–1297, doi:10.1161/CIRCRESAHA.110.233056 (2011).

13 Melki, I., Tessandier, N., Zufferey, A. & Boilard, E. Platelet microvesicles in health and disease. Platelets 28, 214–221, doi:10.1080/09537104.2016.1265924 (2017).

14 Ridger, V. C. et al.. Microvesicles in vascular homeostasis and diseases. Position Paper of the European Society of Cardiology (ESC) Working Group on Atherosclerosis and Vascular Biology. Thromb Haemost 117, 1296–1316, doi:10.1160/TH16-12-0943 (2017).

15 Su, Y. et al.. Multi-Omics Resolves a Sharp Disease-State Shift between Mild and Moderate COVID-19. Cell 183, 1479-1495 e1420, doi:10.1016/j.cell.2020.10.037 (2020).

16 Del Valle, D. M. et al.. An inflammatory cytokine signature predicts COVID-19 severity and survival. Nat Med 26, 1636–1643, doi:10.1038/s41591-020-1051-9 (2020).

17 Zheng, M. et al.. Functional exhaustion of antiviral lymphocytes in COVID-19 patients. Cell Mol Immunol 17, 533–535, doi:10.1038/s41423-020-0402-2 (2020).

18 Cao, X. COVID-19: immunopathology and its implications for therapy. Nat Rev Immunol 20, 269–270, doi:10.1038/s41577-020-0308-3 (2020).

19 Yang, X. et al.. Clinical course and outcomes of critically ill patients with SARS-CoV-2 pneumonia in Wuhan, China: a single-centered, retrospective, observational study. Lancet Respir Med 8, 475–481, doi:10.1016/S2213-2600(20)30079-5 (2020).

20 Huang, C. et al.. Clinical features of patients infected with 2019 novel coronavirus in Wuhan, China. Lancet 395, 497–506, doi:10.1016/S0140-6736(20)30183-5 (2020).

21 Liu, J. et al.. Longitudinal characteristics of lymphocyte responses and cytokine profiles in the peripheral blood of SARS-CoV-2 infected patients. EBioMedicine 55, 102763, doi:10.1016/j.ebiom.2020.102763 (2020).

22 Chen, R. et al.. Longitudinal hematologic and immunologic variations associated with the progression of COVID-19 patients in China. J Allergy Clin Immunol 146, 89–100, doi:10.1016/j.jaci.2020.05.003 (2020).

23 Mathew, D. et al.. Deep immune profiling of COVID-19 patients reveals distinct immunotypes with therapeutic implications. Science 369, doi:10.1126/science.abc8511 (2020).

24 Laing, A. G. et al.. A dynamic COVID-19 immune signature includes associations with poor prognosis. Nat Med 26, 1623–1635, doi:10.1038/s41591-020-1038-6 (2020).

25 Hotchkiss, R. S. & Nicholson, D. W. Apoptosis and caspases regulate death and inflammation in sepsis. Nat Rev Immunol 6, 813–822, doi:10.1038/nri1943 (2006).

26 Liao, M. et al.. Single-cell landscape of bronchoalveolar immune cells in patients with COVID-19. Nat Med 26, 842–844, doi:10.1038/s41591-020-0901-9 (2020).

27 Rydyznski Moderbacher, C. et al. Antigen-Specific Adaptive Immunity to SARS-CoV-2 in Acute COVID-19 and Associations with Age and Disease Severity. Cell 183, 996–1012 e1019, doi:10.1016/j.cell.2020.09.038 (2020).

28 Sekine, T. et al.. Robust T Cell Immunity in Convalescent Individuals with Asymptomatic or Mild COVID-19. Cell 183, 158–168 e114, doi:10.1016/j.cell.2020.08.017 (2020).

29 Zhou, R. et al.. Acute SARS-CoV-2 Infection Impairs Dendritic Cell and T Cell Responses. Immunity 53, 864–877 e865, doi:10.1016/j.immuni.2020.07.026 (2020).

30 Sette, A. & Crotty, S. Adaptive immunity to SARS-CoV-2 and COVID-19. Cell 184, 861–880, doi:10.1016/j.cell.2021.01.007 (2021).

31 Braun, J. et al.. SARS-CoV-2-reactive T cells in healthy donors and patients with COVID-19. Nature 587, 270–274, doi:10.1038/s41586-020-2598-9 (2020).

32 Tan, A. T. et al.. Early induction of SARS-CoV-2 specific T cells associates with rapid viral clearance and mild disease in COVID-19 patient . sbioRxiv, 1-26, doi:10.1101/2020.10.15.341958 (2020).

33 Kranich, J. et al.. In vivo identification of apoptotic and extracellular vesicle-bound live cells using image-based deep learning. J Extracell Vesicles 9, 1792683, doi:10.1080/20013078.2020.1792683 (2020).

34 Trautz, B. et al.. The host-cell restriction factor SERINC5 restricts HIV-1 infectivity without altering the lipid composition and organization of viral particles. J Biol Chem 292, 13702–13713, doi:10.1074/jbc.M117.797332 (2017).

35 Li, Q. et al.. Eosinopenia and elevated C-reactive protein facilitate triage of COVID-19 patients in fever clinic: A retrospective case-control study. EClinicalMedicine 23, 100375, doi:10.1016/j.eclinm.2020.100375 (2020).

36 Lind, S. E. Phosphatidylserine is an overlooked mediator of COVID-19 thromboinflammation. Heliyon 7, e06033, doi:10.1016/j.heliyon.2021.e06033 (2021).

37 WorldHealthOrganization. Novel Coronavirus: COVID-19 Therapeutic Trial Synopsis. . WHO R&D Blueprint Draft February 18, 2020. www.who.int/blueprint/priority-diseases/key-action/COVID-19_Treatment_Trial_Design_Master_Protocol_synopsis_Final_ 18022020.pdf (2020).

38 Connor, D. E., Exner, T., Ma, D. D. & Joseph, J. E. The majority of circulating platelet-derived microparticles fail to bind annexin V, lack phospholipid-dependent procoagulant activity and demonstrate greater expression of glycoprotein Ib. Thromb Haemost 103, 1044–1052, doi:10.1160/TH09-09-0644 (2010).

39 Lippi, G. & Plebani, M. Laboratory abnormalities in patients with COVID-2019 infection. Clin Chem Lab Med 58, 1131–1134, doi:10.1515/cclm-2020-0198 (2020).

40 Mathieu, M., Martin Jaular, L., Lavieu, G. & Théry, C. Specificities of secretion and uptake of exosomes and other extracellular vesicles for cell-to-cell communication. Nature cell biology 21, 1–9, doi:papers3://publication/doi/10.1038/s41556-018-0250-9 (2018).

41 Reininger, A. J. et al.. Mechanism of platelet adhesion to von Willebrand factor and microparticle formation under high shear stress. Blood 107, 3537–3545, doi:10.1182/blood-2005-02-0618 (2006).

42 Ponomareva, A. A. et al.. Intracellular origin and ultrastructure of platelet-derived microparticles. J Thromb Haemost 15, 1655–1667, doi:10.1111/jth.13745 (2017).

43 Arraud, N. et al.. Extracellular vesicles from blood plasma: determination of their morphology, size, phenotype and concentration. Journal of Thrombosis and Haemostasis 12, 614–627, doi:papers3://publication/doi/10.1111/jth.12554 (2014).

44 Teuwen, L. A., Geldhof, V., Pasut, A. & Carmeliet, P. COVID-19: the vasculature unleashed. Nat Rev Immunol 20, 389–391, doi:10.1038/s41577-020-0343-0 (2020).

45 Klok, F. A. et al.. Incidence of thrombotic complications in critically ill ICU patients with COVID-19. Thromb Res 191, 145–147, doi:10.1016/j.thromres.2020.04.013 (2020).

46 Tang, N., Li, D., Wang, X. & Sun, Z. Abnormal coagulation parameters are associated with poor prognosis in patients with novel coronavirus pneumonia. J Thromb Haemost 18, 844–847, doi:10.1111/jth.14768 (2020).

47 Middeldorp, S. et al.. Incidence of venous thromboembolism in hospitalized patients with COVID-19. J Thromb Haemost 18, 1995–2002, doi:10.1111/jth.14888 (2020).

48 Heijnen, H. F., Schiel, A. E., Fijnheer, R., Geuze, H. J. & Sixma, J. J. Activated platelets release two types of membrane vesicles: microvesicles by surface shedding and exosomes derived from exocytosis of multivesicular bodies and alpha-granules. Blood 94, 3791–3799 (1999).

49 Aatonen, M. T. et al.. Isolation and characterization of platelet-derived extracellular vesicles. J Extracell Vesicles 3, doi:10.3402/jev.v3.24692 (2014).

50 Zaid, Y. et al.. Platelets Can Associate with SARS-Cov-2 RNA and Are Hyperactivated in COVID-19. Circ Res, doi:10.1161/CIRCRESAHA.120.317703 (2020).

51 Manne, B. K. et al.. Platelet gene expression and function in patients with COVID-19. Blood 136, 1317–1329, doi:10.1182/blood.2020007214 (2020).

52 van der Zee, P. M. et al.. P-selectin-and CD63-exposing platelet microparticles reflect platelet activation in peripheral arterial disease and myocardial infarction. Clin Chem 52, 657–664, doi:10.1373/clinchem.2005.057414 (2006).

53 Rolfes, V. et al.. PD-L1 is expressed on human platelets and is affected by immune checkpoint therapy. Oncotarget 9, 27460–27470, doi:10.18632/oncotarget.25446 (2018).

54 Chapman, L. M. et al.. Platelets present antigen in the context of MHC class I. J Immunol 189, 916–923, doi:10.4049/jimmunol.1200580 (2012).

55 DeGregori, J. & Johnson, D. G. Distinct and Overlapping Roles for E2F Family Members in Transcription, Proliferation and Apoptosis. Curr Mol Med 6, 739–748, doi:10.2174/1566524010606070739 (2006).

56 Song, J., So, T., Cheng, M., Tang, X. & Croft, M. Sustained survivin expression from OX40 costimulatory signals drives T cell clonal expansion. Immunity 22, 621–631, doi:10.1016/j.immuni.2005.03.012 (2005).

57 Sachs, A. B. Cell cycle-dependent translation initiation: IRES elements prevail. Cell 101, 243–245, doi:10.1016/s0092-8674(00)80834-x (2000).

58 Scholzen, T. & Gerdes, J. The Ki-67 protein: From the known and the unknown. Journal of Cellular Physiology 182, 311–322, doi:10.1002/(sici)1097-4652(200003)182:3<311::Aid-jcp1>3.0.Co;2-9 (2000).

59 Kared, H. et al.. SARS-CoV-2-specific CD8+ T cell responses in convalescent COVID-19 individuals. J Clin Invest 131, doi:10.1172/JCI145476 (2021).

60 Best, J. A. et al.. Transcriptional insights into the CD8+ T cell response to infection and memory T cell formation. Nature Immunology 14, 404–412, doi:papers3://publication/doi/10.1038/ni.2536 (2013).

61 Sallusto, F., Geginat, J. & Lanzavecchia, A. Central memory and effector memory T cell subsets: function, generation, and maintenance. Annu Rev Immunol 22, 745–763, doi:10.1146/annurev.immunol.22.012703.104702 (2004).

62 Cappellano, G. et al.. Circulating Platelet-Derived Extracellular Vesicles Are a Hallmark of Sars-Cov-2 Infection. Cells 10, doi:10.3390/cells10010085 (2021).

63 Mazzoni, A. et al.. Impaired immune cell cytotoxicity in severe COVID-19 is IL-6 dependent. J Clin Invest 130, 4694–4703, doi:10.1172/JCI138554 (2020).

64 Song, W. C. & FitzGerald, G. A. COVID-19, microangiopathy, hemostatic activation, and complement. J Clin Invest 130, 3950–3953, doi:10.1172/JCI140183 (2020).

65 Sims, P. J., Faioni, E. M., Wiedmer, T. & Shattil, S. J. Complement proteins C5b-9 cause release of membrane vesicles from the platelet surface that are enriched in the membrane receptor for coagulation factor Va and express prothrombinase activity. J Biol Chem 263, 18205–18212 (1988).

66 Mahase, E. Long covid could be four different syndromes, review suggests. BMJ 371, m3981, doi:10.1136/bmj.m3981 (2020).

67 Zheng, H. Y. et al.. Elevated exhaustion levels and reduced functional diversity of T cells in peripheral blood may predict severe progression in COVID-19 patients. Cell Mol Immunol 17, 541–543, doi:10.1038/s41423-020-0401-3 (2020).

68 Green, S. A. et al.. Activated platelet-T-cell conjugates in peripheral blood of patients with HIV infection: coupling coagulation/inflammation and T cells. AIDS 29, 1297–1308, doi:10.1097/QAD.0000000000000701 (2015).

69 Polasky, C., Wendt, F., Pries, R. & Wollenberg, B. Platelet Induced Functional Alteration of CD4(+) and CD8(+) T Cells in HNSCC. International journal of molecular sciences 21, doi:10.3390/ijms21207507 (2020).

70 De Biasi, S. et al.. Marked T cell activation, senescence, exhaustion and skewing towards TH17 in patients with COVID-19 pneumonia. Nat Commun 11, 3434, doi:10.1038/s41467-020-17292-4 (2020).

71 Song, J. W. et al.. Immunological and inflammatory profiles in mild and severe cases of COVID-19. Nat Commun 11, 3410, doi:10.1038/s41467-020-17240-2 (2020).

72 Diao, B. et al.. Reduction and Functional Exhaustion of T Cells in Patients With Coronavirus Disease 2019 (COVID-19). Front Immunol 11, 827, doi:10.3389/fimmu.2020.00827 (2020).

73 Chen, G. et al.. Exosomal PD-L1 contributes to immunosuppression and is associated with anti-PD-1 response. Nature 560, 382–386, doi:10.1038/s41586-018-0392-8 (2018).

74 Rha, M. S. et al.. PD-1-Expressing SARS-CoV-2-Specific CD8(+) T Cells Are Not Exhausted, but Functional in Patients with COVID-19. Immunity 54, 44–52 e43, doi:10.1016/j.immuni.2020.12.002 (2021).

75 Hottz, E. D. et al.. Platelet activation and platelet-monocyte aggregate formation trigger tissue factor expression in patients with severe COVID-19. Blood 136, 1330–1341, doi:10.1182/blood.2020007252 (2020).

