## Supplementary files for "Binding of phosphatidylserine-positive microparticles by PBMCs classifies disease severity in COVID-19 patients"

**TITLE**

**AFFILIATIONS**

Großhaderner Str. 9, D-82152 Planegg-Martinsried, Germany

| | COVID-ALL | mild | moderate | severe |
|---|---|---|---|---|
| **Parameter** | **Value** | **Value** | **Value** | **Value** |
| **Demographic data** | | | | |
| Number | 54 | 17 | 22 | 15 |
| Age (y) | 63 (31-89) | 55 (23-80) | 62 (32-89) | 55 (23-80) |
| Gender (f/m) | 18 (33%)/36(67%) | 6 (35%)/11(65%) | 6 (27%)/16 (73%) | 6 (40%)/9(60%) |
| **Medical history** | | | | |
| Hypertension | 21 (39%) | 2 (12%) | 11 (50%) | 8 (53%) |
| Diabetes mellitus | 15 (28%) | 3 (18%) | 6 (27%) | 6 (40%) |
| Coronary artery disease | 11 (20%) | 3 (18%) | 4 (18%) | 4 (27%) |
| Myocardial infarction | 5 (9%) | 2 (12%) | 1 (5%) | 2 (13%) |
| Atrial fibrillation | 11 (20%) | 2 (12%) | 6 (27%) | 3 (20%) |
| Congenital heart failure | 5 (9%) | 2 (12%) | 3 (14%) | 0 |
| Peripheral artery disease | 3 (6%) | 2 (12%) | 1 (5%) | 0 |
| Dementia | 4 (7%) | 1 (6%) | 1 (5%) | 2 (13%) |
| Stroke | 6 (11%) | 3 (18%) | 1 (5%) | 2 (13%) |
| Chronic obstructive pulmonary disease | 4 (7%) | 1 (6%) | 1 (5%) | 2 (13%) |
| Asthma | 4 (7%) | 2 (12%) | 1 (5%) | 1 (7%) |
| Lymphoma | 1 (2%) | 0 | 1 (5%) | 0 |
| Solid Tumors | 8 (15%) | 1 (6%) | 4 (18%) | 3 (20%) |
| Chemotherapy | 2 (4%) | 1 (6%) | 1 (5%) | 0 |
| Chronic kidney disease | 10 (19%) | 3 (18%) | 4 (18%) | 3 (20%) |
| Transplantation | 4 (7%) | 0 | 4 (18%) | 0 |
| Rheumatic disease | 1 (2%) | 0 | 0 | 1 (7%) |
| Active smoker | 2 (4%) | 1 (6%) | 0 | 1 (7%) |
| **Medication at admission** | | | | |
| Immunsuppression | 10 (19%) | 2 (12%) | 7 (32%) | 1 (7%) |
| Aspirin | 12 (22%) | 4 (24%) | 3 (14%) | 5 (33%) |
| **Disease characteristics** | | | | |
| WHO deterioration | 36 (67%) | 4 (24%) | 17 (77%) | 15 (100%) |
| Number of days since symtoms started | 22 (3-61) | 17 (5-48) | 19 (3-51) | 29 (5-61) |
| Number of days since first diagnosis | 20 (2-76) | 19 (2-48) | 20 (3-76) | 21 (2-58) |
| Days of Hospitalization | 19 (3-67/ongoing) | 11 (3-46) | 21 (5-56) | 26 (5-67/ongoing) |
| WHOmax | 4 (1-8) | 3 (1-3) | 4 | 6 (5-8) |
| Mortality | 6 (11%) | 0 | 0 | 6 (40%) |
| ECMO | 2 (4%) | 0 | 0 | 2 (13%) |
| Non-invasive Ventilation | 6 (11%) | 1 | 0 | 6 (40%) |
| Invasive Ventilation | 7 (13%) | 0 | 0 | 7 (47%) |
| Mechanical ventilation (days) | 15 (3-61/ongoing) | ongoing | 0 | 15 (3-61) |
| **Disease treatment** | | | | |
| Hydroxychloroquine | 1 (2%) | 0 | 0 | 1 (7%) |
| Glucocorticoids | 35 (65%) | 7 (41%) | 17 (77%) | 11 (73%) |
| Remdesevir | 9 (17%) | 3 (18%) | 5 (23%) | 1 (8%) |
| Treatment by convalescent plasma | 1 (2%) | 0 | 0 | 1 (7%) |
| Tocilizumab | 2 (4%) | 0 | 0 | 2 (13%) |
| Cytosorb | 2 (4%) | 0 | 0 | 2 (13%) |
| ACE inhibitors | 5 (9%) | 5 (29%) | 0 | 0 |
| Ruxolitinib | 2 (4%) | 0 | 1 (5%) | 1 (7%) |
| **Lab values** | | | | |
| nearest CRP | 4.1 (0.1-17.6) | 2.8 (0.1-12.8) | 3.0 (0.3-8.6) | 6.8 (0.1-17.6) |
| nearest D-dimer | 2.0 (0.2-12.7) | 2.5 (0.2-12.7) | 1.7 (0.3-6.3) | 2.0 (0.5-7.0) |
| nearest Ferritin | 1173 (33-10110) | 1524 (34-10110) | 921 (115-3047) | 1184 (33-6570) |
| nearest Fibrinogen | 466 (123.5-900) | 429 (204-900) | 474 (305-641) | 483.5 (123.5-778) |
| nearest IL6 | 42 (1.5-226) | 22.2 (1.5-83.2) | 25.2 (2.0-123) | 84.5 (2.5-226) |
| nearest INR | 1.1 (0.8-3.4) | 1.1 (0.9-1.9) | 1.2 (0.8-3.4) | 1.1 (0.9-1.3) |
| nearest Leukocytes | 7.5 (1.21-27.2) | 6.4 (2.1-11.5) | 6.9 (1.2-15.7) | 9.4 (3.2-27.2.5) |
| nearest Lymphocytes (%) | 21.1 (2-59) | 27.5 (15-59) | 21.6 (5-47) | 13.8 (2-59) |
| nearest Lymphocytes (G/L) | 1.2 (0.3-3.0) | 1.5 (0.7-2.5) | 1.2 (0.3-2.5) | 1.0 (0.3-3.0) |
| nearest Monocytes (%) | 8.1 (1-29) | 8.5 (4-16) | 9.3 (3-29) | 5.8 (4-14) |
| nearest Monocytes (G/L) | 0.6 (0.1-1.5) | 0.6 (0.1-1.1) | 0.6 (0.1-1.5) | 0.6 (0.3-1.1) |
| nearest Neutrophils (%) | 68.4 (43-93) | 62.8 (46-75) | 66.3 (43-89) | 77.2 (49.5-93) |
| nearest Neutrophils (G/L) | 5.2 (0.5-25.2) | 4.0 (0.8-8.7) | 4.4 (0.5-11.9) | 7.4 (2.0-25.2) |
| nearest PCT | 0.6 (0.1-7.9) | 0.6 (0.1-7.9) | 0.2 (0.1-0.5) | 1.1 (0.1-6.9) |
| nearest Platelets | 259 (15-585) | 283 (15-585) | 252 (20-554) | 247 (38-469) |
| nearest PTT | 33.0 (20-81) | 29.6 (21-56.5) | 32.8 (23-57) | 36.5 (20-81) |
| CRP at admission | 5.6 (0.1-20.1) | 4.6 (0.1-20.1) | 3.8 (0.4-9.9) | 9.2 (0.9-16.8) |
| D-dimer at admission | 3.1 (0.2-20.9) | 3.9 (0.2-19.5) | 2.3 (0.3-8.6) | 3.6 (0.5-20.9) |
| Ferritin at admission | 988 (63-7962) | 1592 (63-7962) | 548 (115-1118) | 962 (77-3577) |
| Fibrinogen at admission | 493.6 (204-760) | 384.8 (204-536) | 479.1 (297-695) | 575.9 (469-760) |
| IL6 at admission | 50.3 (2.2-173) | 35.3 (2.2-105) | 44.9 (5.2-138) | 71.7 (21.5-173) |
| INR at admission | 1.1 (0.8-2.5) | 1.0 (0.8-1.9) | 1.1 (0.8-2.5) | 1.0 (0.8-1.2) |
| Leukocytes at admission | 6.6 (1.4-23.6) | 5.6 (1.4-11) | 5.4 (1.8-11) | 9.1 (3.1-23.6) |
| Lymphocytes (%) at admission | 19.4 (1-59) | 25.7 (5-59) | 20.8 (4-45) | 11.1 (1-24) |
| Lymphocytes (G/L) at admission | 1.1 (0.2-3.2) | 1.4 (0.7-3.2) | 1.1 (0.2-2.9) | 0.8 (0.2-1.4) |
| Monocytes (%) at admission | 8.4 (2-42) | 8.7 (2-23) | 9.8 (3-42) | 6.1 (2-13) |
| Monocytes (G/L) at admission | 0.5 (0.1-1.5) | 0.5 (0.1-1.0) | 0.5 (0.1-1.5) | 0.5 (0.2-1.1) |
| Neutrophils (%) at admission | 69.2 (39-92) | 62.6 (39-80) | 67.4 (45-92) | 78.8 (64-91) |
| Neutrophils (G/L) at admission | 4.6 (0.6-22.1) | 3.4 (0.8-7.3) | 0.5 (0.6-10.1) | 7.2 (2.1-22.1) |
| PCT at admission | 0.8 (0.1-7.9) | 0.8 (0.1-7.9) | 0.2 (0.1-0.5) | 1.1 (0.1-5.1) |
| Platelets at admission | 225.6 (39-489) | 237.6 (39-463) | 210.6 (82-350) | 233.5 (133-489) |
| PTT at admission | 29.3 (19-56) | 28.9 (19-56) | 31.0 (20-45) | 27.4 (23-37) |
| CRP last value | 3.5 (0.1-18.8) | 3.0 (0.1-12.3) | 2.1 (0.1-7.6) | 5.5 (0.1-18.8) |
| D-dimer last value | 2.7 (0.2-16.3) | 2.8 (0.2-12.7) | 1.8 (0.3-6.3) | 3.5 (0.5-16.3) |
| Ferritin last value | 988 (25-10110) | 1364 (25-10110) | 645 (117-1986) | 1144 (33-6570) |
| Fibrinogen last value | 423.0 (138-666) | 399.8 (204-536) | 435.7 (305-666) | 425.5 (138-634) |
| IL6 last value | 58.3 (0.9-910) | 29.7 (0.9-185) | 18.0 (2.7-78) | 129.6 (2.5-910) |
| INR last value | 1.1 (0.8-2.8) | 1.1 (0.9-2.3) | 1.1 (0.8-2.8) | 1.1 (0.9-1.9) |
| Leukocytes last value | 8.8 (1.6-31.5) | 6.6 (1.6-11.6) | 9.0 (2.6-20.5) | 10.4 (3.2-31.5) |
| Lymphocytes (%) last value | 20.1 (2-59) | 26.1 (15-59) | 20.4 (7-39) | 14.2 (2-46) |
| Lymphocytes (G/L) last value | 1.6 (0.3-7.7) | 1.6 (0.6-2.7) | 1.8 (0.5-7.7) | 1.2 (0.3-3.3) |
| Monocytes (%) last value | 8.7 (2-20) | 8.6 (5-13) | 9.5 (2-20) | 7.8 (2-14) |
| Monocytes (G/L) last value | 0.7 (0.1-1.7) | 0.6 (0.1-1.1) | 0.8 (0.2-1.7) | 0.6 (0.3-1.2) |
| Neutrophils (%) last value | 66.4 (47-93) | 63.2 (47-49) | 64.2 (48-88) | 73.2 (47-93) |
| Neutrophils (G/L) last value | 5.9 (0.8-29.3) | 4.2 (0.8-7.9) | 5.5 (1.3-11.9) | 8.1 (2.0-29.3) |
| PCT last value | 1.0 (0.1-7.9) | 1.5 (0.1-7.9) | 0.2 (0.1-0.2) | 1.6 (0.1-4.8) |
| Platelets last value | 290.8 (15-624) | 302.4 (15-585) | 293 (86-624) | 275.9 (31-483) |
| PTT last value | 30.4 (22-93) | 29.2 (22-67) | 26.6 (22-32) | 36.8 (22-93) |
| CRP min | 1.5 (0.1-12.3) | 1.5 (0.1-12.3) | 0.6 (0.1-2.5) | 2.5 (0.1-10.9) |
| D-dimer min | 1.4 (0.2-6.3) | 1.5 (0.3-3.5) | 1.6 (0.3-3.6) | 1.0 (0.2-2.4) |
| Ferritin min | 960 (25-9297) | 1495.5 (25-9297) | 746.1 (113-3047) | 776.9 (33-3577) |
| Fibrinogen min | 386.8 (97-653) | 352.7 (204-536) | 402.6 (297-499) | 392.4 (97-653) |
| IL6 min | 14.1 (0.9-107) | 14.1 (1.6-61) | 7.8 (1.5-35.8) | 22.5 (2.5-107) |
| INR min | 0.9 (0.8-2.5) | 1.0 (0.8-1.7) | 1.0 (0.8-2.5) | 0.9 (0.8-1.1) |
| Leukocytes min | 4.5 (0.6-13.5) | 4.5 (1.0-7.8) | 3.9 (1.0-8.1) | 5.1 (0.6-13.9) |
| Lymphocytes (%) min | 12.3 (1-59) | 19.3 (5-59) | 13.2 (4-35) | 4.7 (1-18) |
| Lymphocytes (G/L) min | 0.9 (0.2-2.2) | 1.3 (0.6-2.2) | 0.8 (0.3-1.2) | 0.6 (0.2-1.4) |
| Monocytes (%) min | 4.6 (1-13) | 6.2 (1-12) | 4.8 (1-13) | 2.7 (1-9) |
| Monocytes (G/L) min | 0.4 (0.1-1.9) | 0.5 (0.1-0.9) | 0.3 (0.1-0.9) | 0.4 (0.1-0.7) |
| Neutrophils (%) min | 60.2 (39-90) | 56.5 (39-79) | 57.4 (40-84) | 68.4 (47-90) |
| Neutrophils (G/L) min | 2.9 (0.1-12.6) | 2.7 (0.1-5.0) | 2.5 (0.4-5.0) | 3.6 (0.1-12.6) |
| PCT min | 0.5 (0.1-7.9) | 1.4 (0.1-7.9) | 0.1 (0.1-0.2) | 0.5 (0.1-3.3) |
| Platelets min | 167.9 (12-370) | 202.3 (15-331) | 151.6 (12-335) | 157.4 (21-370) |
| PTT min | 24.5 (18-54) | 26.8 (19-54) | 24.0 (18-32) | 23 (19-32) |
| CRP max | 10.2 (0.1-34.5) | 8.5 (0.1-22) | 6.8 (1.3-16.3) | 16.2 (1.1-34.5) |
| D-dimer max | 4.4 (0.3-31.1) | 4.6 (0.3-19.5) | 2.0 (0.3-6.5) | 7.3 (1.1-31.1) |
| Ferritin max | 1742.9 (84-17947) | 2533.8 (84-17947) | 1027.7 (236-3047) | 2058 (245-11078) |
| Fibrinogen max | 553.3 (204-883) | 408 (204-536) | 511 (357-695) | 679.4 (506-883) |
| IL6 max | 654.9 (8.1-14510) | 107.7 (8.1-757) | 71.8 (18.9-161) | 1906.5 (48.5-14510) |
| INR max | 1.3 (0.9-3.8) | 1.2 (0.9-2.3) | 1.3 (0.9-3.8) | 1.3 (1.0-2.2) |
| Leukocytes max | 12.4 (2.7-33) | 8.5 (2.7-15.3) | 11.2 (3.1-20.5) | 17.6 (5.6-33) |
| Lymphocytes (%) max | 28.5 (4-59) | 34 (15-59) | 30.2 (8-47) | 21.1 (4-59) |
| Lymphocytes (G/L) max | 1.8 (0.4-7.7) | 1.9 (0.7-3.6) | 2.1 (0.6-7.7) | 1.5 (0.4-3.3) |
| Monocytes (%) max | 12.4 (5-42) | 11.4 (6-23) | 13.8 (7-42) | 11.5 (5-22) |
| Monocytes (G/L) max | 0.9 (0.1-1.9) | 0.7 (0.1-1.2) | 0.9 (0.2-1.7) | 0.9 (0.3-1.5) |
| Neutrophils (%) max | 76.8 (51-94) | 69.5 (51-80) | 76.3 (57-92) | 85 (64-94) |
| Neutrophils (G/L) max | 9.2 (1.7-29.3) | 5.4 (1.7-9.2) | 7.8 (1.7-14.1) | 14.6 (3.1-29.3) |
| PCT max | 2.1 (0.1-29.5) | 1.6 (0.1-7.9) | 0.5 (0.1-3.9) | 4.3 (0.2-29.5) |
| Platelets max | 356.7 (45-676) | 329.7 (45-585) | 343.2 (82-624) | 400.1 (143-676) |
| PTT max | 47.3 (23-156) | 34.9 (23-67) | 41.9 (25-90) | 66.1 (25-156) |

**Supplemental Table 1. COVID-19 patient details.**

| Parameter | Value |
|---|---|
| **Demographic data** | |
| Number | 12 |
| Age (y) | 40 (21-72) |
| Gender (f/m) | 8 (67%)/4(33%) |
| **Disease characteristics** | |
| Number of days since first diagnosis | 182 (69-246) |
| Hospitalized/ambulatory | 4 (33%)/8 (67%) |
| **Disease characteristics hospitalized patients** | |
| WHO deterioration | 3 (75%) |
| Days of Hospitalization | 4.75 (1-11) |
| WHOmax | 2.75 (1-4) |
| Death | 0 |
| ECMO | 0 |
| Non-invasive Ventilation | 0 |
| Invasive Ventilation | 0 |
| Mechanical ventilation (days) | 0 |
| **Medical history hospitalized patients** | |
| Hypertension | 1 (25%) |
| Diabetes mellitus | 0 |
| Coronary artery disease | 0 |
| MyocardialInfarction | 0 |
| AtrialFibrillation | 0 |
| CongHeartFailure | 0 |
| Peripheral artery disease | 0 |
| Dementia | 0 |
| Stroke | 0 |
| Chronic obstructive pulmonary disease | 0 |
| Asthma | 0 |
| Lymphoma | 0 |
| Solid Tumors | 1 (25%) |
| Chemotherapy | 0 |
| chronic kidney disease | 0 |
| Dialysis | 0 |
| Transplantation | 0 |
| Rheumatic disease | 0 |
| active smoker | 0 |

**Supplemental Table 2. Recovered patient details.**

| Parameter | Value |
|---|---|
| **Demographic data** | |
| Number | 35 |
| Age (y) | 39 (22-81) |
| Gender (f/m) | 17 (49%)/18(51%) |

**Supplemental Table 3. Healthy donor details.**

| Vendor | Cat# | Antibody/dye | Clone |
|---|---|---|---|
| BioLegend | 325630 | CD14 AF594 | HCD14 |
| BioLegend | 301854 | CD14 APCFire750 | M5E2 |
| BioLegend | 360706 | CD16 APC | B73.1 |
| BioLegend | 360726 | CD16 APCFire750 | B73.1 |
| BioLegend | 302258 | CD19 APCFire750 | HIB19 |
| BioLegend | 302208 | CD19 PE | HIB19 |
| Invitrogen | 2135078 | CD197 APC | 3D12 |
| BioLegend | 329742 | CD274 AF594 | 29E2A3 |
| BioLegend | 374507 | CD274 BV421 | MIH3 |
| BioLegend | 300412 | CD3 APC | UCHT1 |
| BioLegend | 300470 | CD3 APCFire750 | UCHT1 |
| BioLegend | 300434 | CD3 BV421 | UCHT1 |
| BioLegend | 300420 | CD3 PECy7 | UCHT1 |
| BioLegend | 300408 | CD3 PE | UCHT1 |
| BioLegend | 303506 | CD38 PE | HIT2 |
| eBioscience | 4272496 | CD4 APC | RPA-T4 |
| BioLegend | 300521 | CD4 PacBlue | RPA-T4 |
| BioLegend | 300512 | CD4 PECy7 | RPA-T4 |
| BioLegend | 303709 | CD41 APC | HIP8 |
| BioLegend | 303730 | CD41 BV421 | HIP8 |
| BioLegend | 304122 | CD45RA PerCPCy5.5 | HI100 |
| BioLegend | 362554 | CD56 APCFire750 | 5.1H11 |
| BioLegend | 304906 | CD62P PE | AK4 |
| BioLegend | 353012 | CD63 Pacific Blue | H5C6 |
| BioLegend | 353003 | CD63 PE | H5C6 |
| BioLegend | 301056 | CD8 AF594 | RPA-T4 |
| BioLegend | 301046 | CD8 BV785 | RPA-T4 |
| BioLegend | 301012 | CD8 PECy7 | RPA-T4 |
| BioLegend | 374211 | CD86 BV421 | BU63 |
| Miltenyi | 5171116501 | FcR Blocking Reagent human | |
| ThermoFisher Scientific | 65-0865-18 | Fixabile Viability Dye eFluor780 | |
| BioLegend | 307616 | HLA-DR PECy7 | L243 |
| BioLegend | 350506 | Ki67 BV421 | Ki-67 |
| self made | | MFG-E8-eGFP (Kranich et al. 2020) | |
| BioLegend | t.b.a. | mC1-biotin | |
| BioLegend | 405237 | Streptavidin AF647 | |
| Aurion | Str-91104/1 | Immuno gold-Streptavidin 6nm | |

Kranich et al 2020 J Extracell Vesicles 2020 Vol. 9 Issue 1 Pages 1792683
DOI: 10.1080/20013078.2020.1792683

**Supplemental Table 4. Antibodies and staining reagents.**

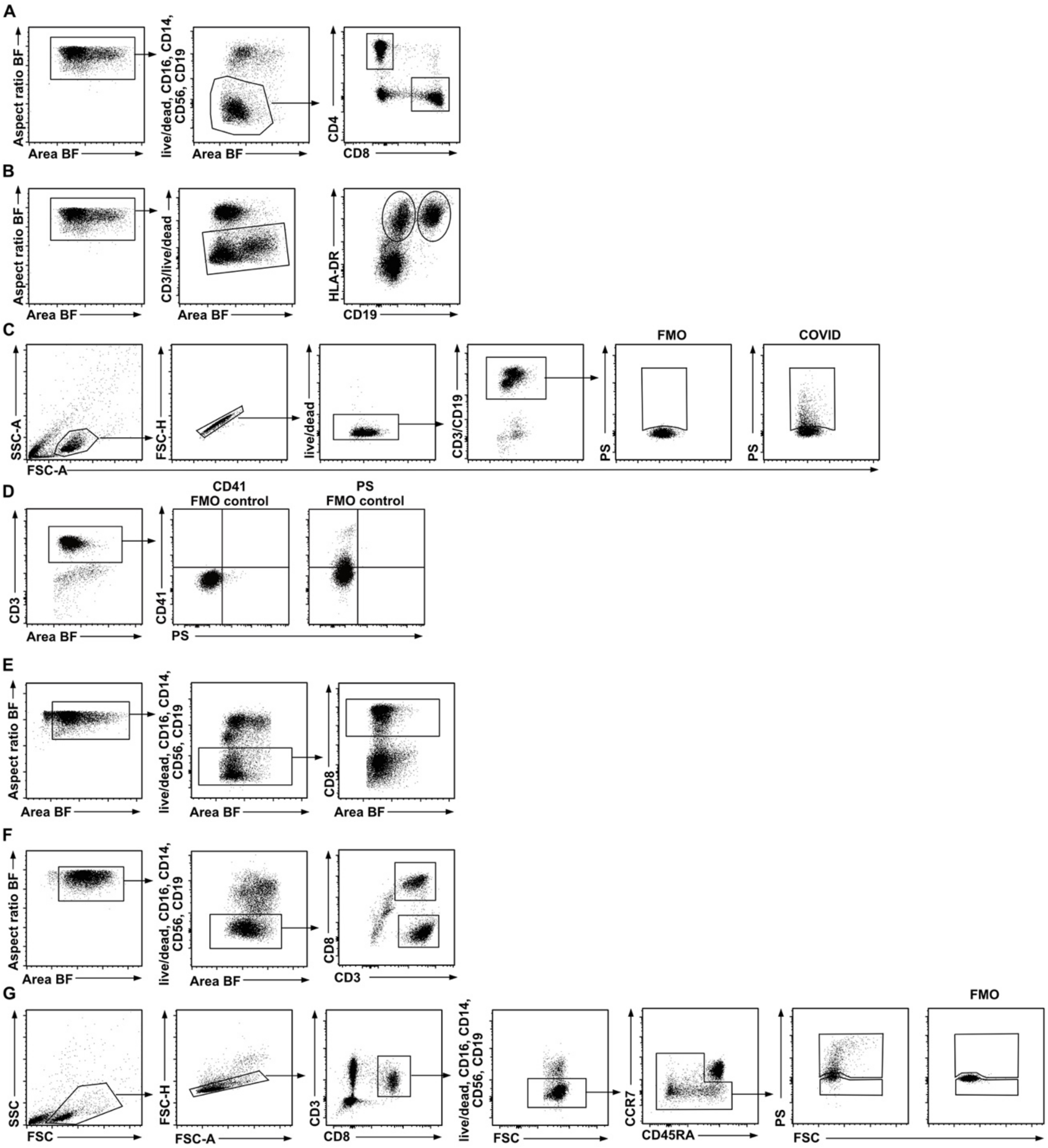


**Suppl. Fig. 1 Flow cytometry gating strategies.** Gating strategy used to examine $CD4^+$ T cells, $CD8^+$ T cells (A) and $HLA\text{-}DR^+CD19^+$ (B cells), $HLA\text{-}DR^+CD19^-$ (non-B/non-T cells) (B) in human PBMCs by flow cytometry. (C) shows the gating strategy used to sort live/$dead^-Tet^+CD3^+$ and $CD19^+$ PBMCs for the analysis and characterization of EVs by TEM (Fig. 4 A-F). (D) Gating strategy used to analyze the presence of CD41 on live/$dead^-CD3^+$ T cells. Representative plots of CD41 and PS FMO controls are shown. (E) Gating strategy for the expression of platelet-markers CD41, CD63, CD62P, and CD274 on live/$dead^-CD8^+$ T cells as shown in Fig. 5. First, single cells were gated using the aspect ratio and the area of the BF channel. Then necrotic live/$dead^+$, $CD19^+$ B cells, $CD56^+$, $CD16^+$, and $CD14^+$ cells were excluded and $CD8^+$ cells were gated for further analysis. (F) Gating strategy for $CD4^+$ and $CD8^+$ T cells used to examine proliferating cells by Ki-67 staining. Cells were first gated on single cells using the aspect ratio and the area of the BF channel. Necrotic live/$dead^+$, $CD19^+$ B cells, $CD56^+$, $CD16^+$, and $CD14^+$ cells were excluded from the analysis and $CD3^+CD8^+$ and $CD3^+CD8^-$ were analyzed for Ki-67 expression as shown in Fig. 6 D,E and Fig. Suppl. 6. (G) Gating strategy used to sort $PS^+$ and $PS^-$ non-naïve $CD8^+$ T cells from PBMCs for RNAseq analysis. First, lymphocytes were gated using the FSC and SSC properties, then single, live/$dead^-$ $CD3^+CD8^+$ T cells were gated. For sorting of non-naïve ($CCR7^{+}CD45RA^{-}$, $CCR7^{-}CD45RA^{-}$ and $CCR7^{-}CD45RA^{+}$) $CD8^{+}$ T cells, $CCR7^{+}CDRA^{+}$ naïve $CD8^{+}$ T cells were excluded. $PS^{+}$ $CD8^{+}$ T cells were gated using FMO controls.

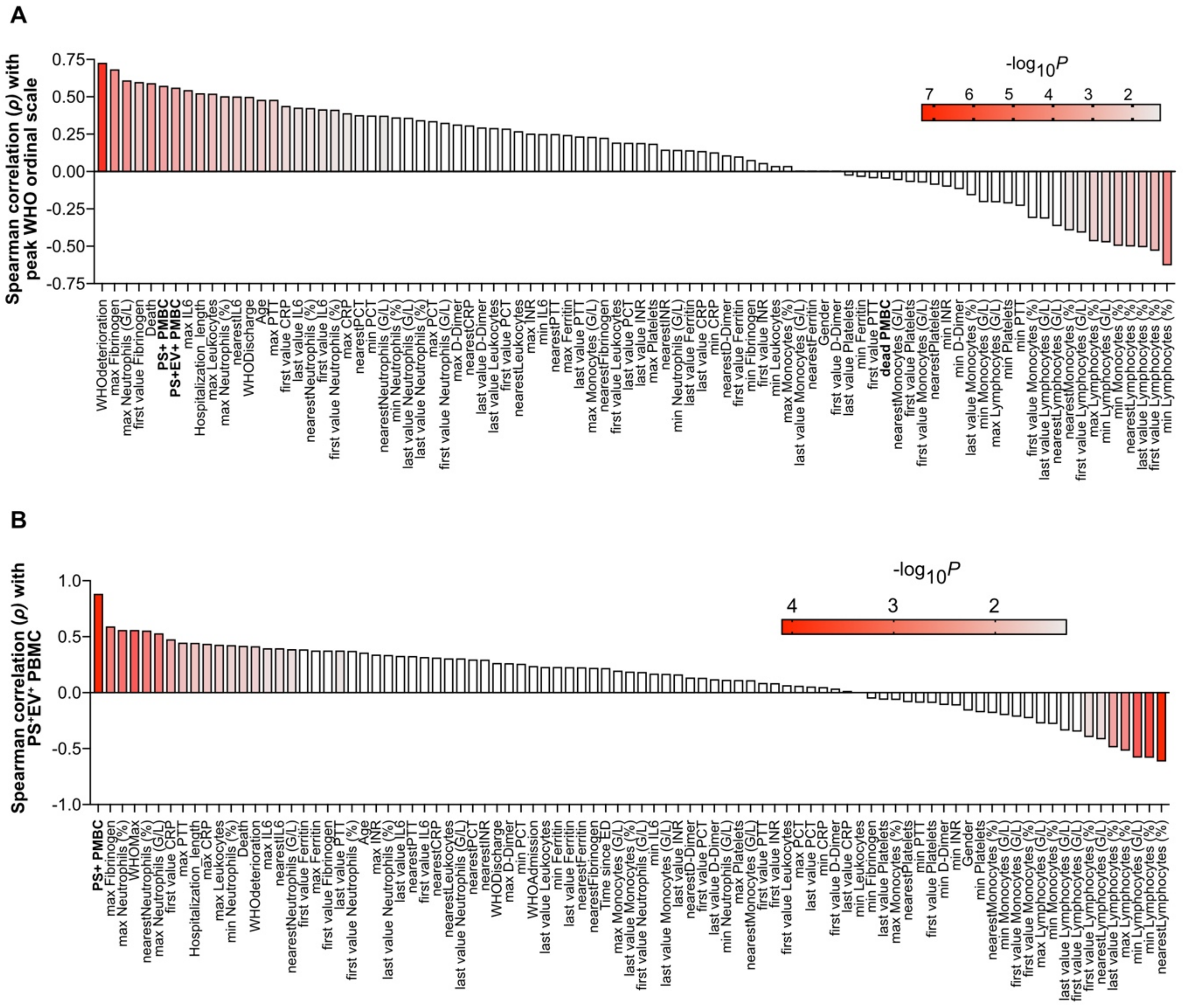


**Suppl. Fig. 2 Summary of spearman correlation of demographic, clinical and immunological features of COVID-19 patients.** Spearman correlation analysis between demographic, clinical and immunological parameters (n= 23-49) and peak WHO ordinal scale (A) or $PS^+EV^+$PBMCs (B) (n=49). Bars display the Spearman's rank correlation coefficients (ρ). P values of indicated clinical and laboratory parameters are indicated by color scale.

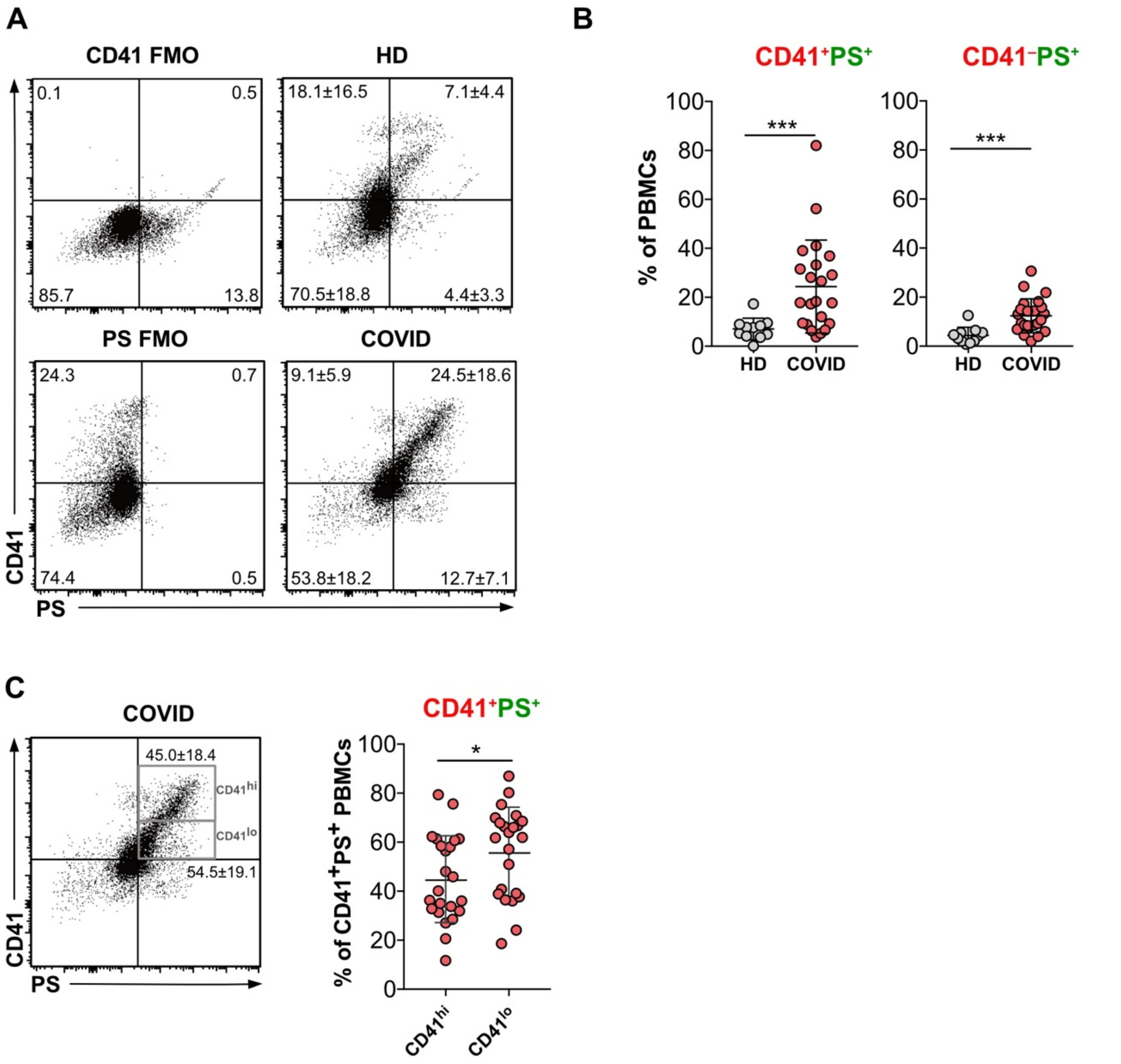


**Suppl. Fig. 3 Analysis of the platelet marker CD41 on PBMCs from COVID-19 patients and healthy donors.** PBMCs from COVID-19 (n=23) and healthy donors (n=11) were analyzed for PS and CD41 positivity. (A) CD41 and PS positive cells were gated using FMO controls. Representative dot plots show the mean frequencies ± SD of the CD41$^{+}$PS$^{-}$, CD41$^{+}$PS$^{+}$, CD41$^{-}$PS$^{+}$, and CD41$^{-}$PS$^{-}$ PBMCs in the respective quadrant. (B) Graphs show the frequencies ± SD of CD41$^{+}$PS$^{+}$ and CD41$^{-}$PS$^{+}$ PBMCs from COVID-19 and healthy donors. (C) Representative dot plot displays the gating and mean frequencies ± SD of PS$^{+}$ CD41$^{hi}$ and PS$^{+}$CD41$^{low}$ PBMCs. Bar graph shows the frequencies ± SD of PS$^{+}$CD41$^{hi}$ and PS$^{+}$CD41$^{low}$ PBMCs in COVID-19 patients. Statistical significance was determined by paired Wilcoxon test and is indicated by asterisks (ns P > 0.5; *P ≤ 0.05; **P ≤ 0.01; ***P ≤ 0.001).

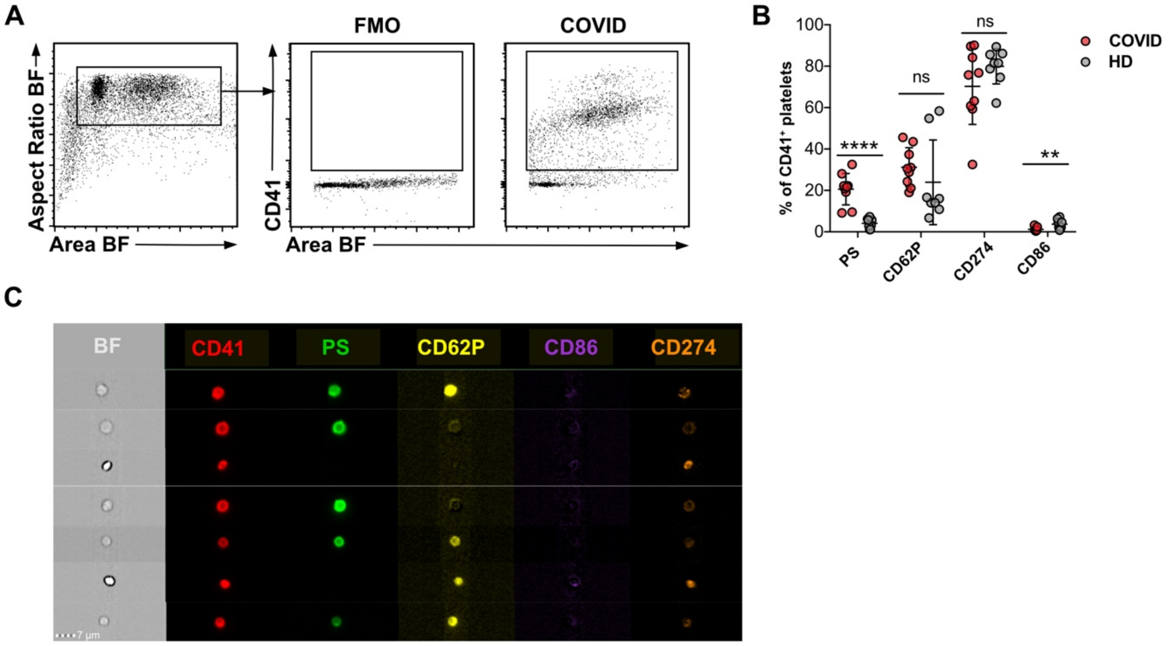


**Suppl. Fig. 4 SARS-COV-2 infections causes platelet activation.** (A) Dot plots illustrate gating of platelets based on aspect ratio and area of the BF channel. Platelets were identified by CD41 expression. (B) Graph shows the frequencies ± SD of PS positivity and of platelet marker (CD62P, CD274, CD86) expression of CD41$^{+}$platelets measured in whole blood by IFC. N = 8 for healthy donor, n = 9 for COVID-19 patient. (C) Representative images of CD41$^{+}$platelets with the platelet markers analyzed from COVID-19 patients. Statistical significance was determined by Mann-Whitney test and is indicated by asterisks ns P > 0.5; *P ≤ 0.05; **P ≤ 0.01; ***P ≤ 0.001; two-tailed unpaired t-test).

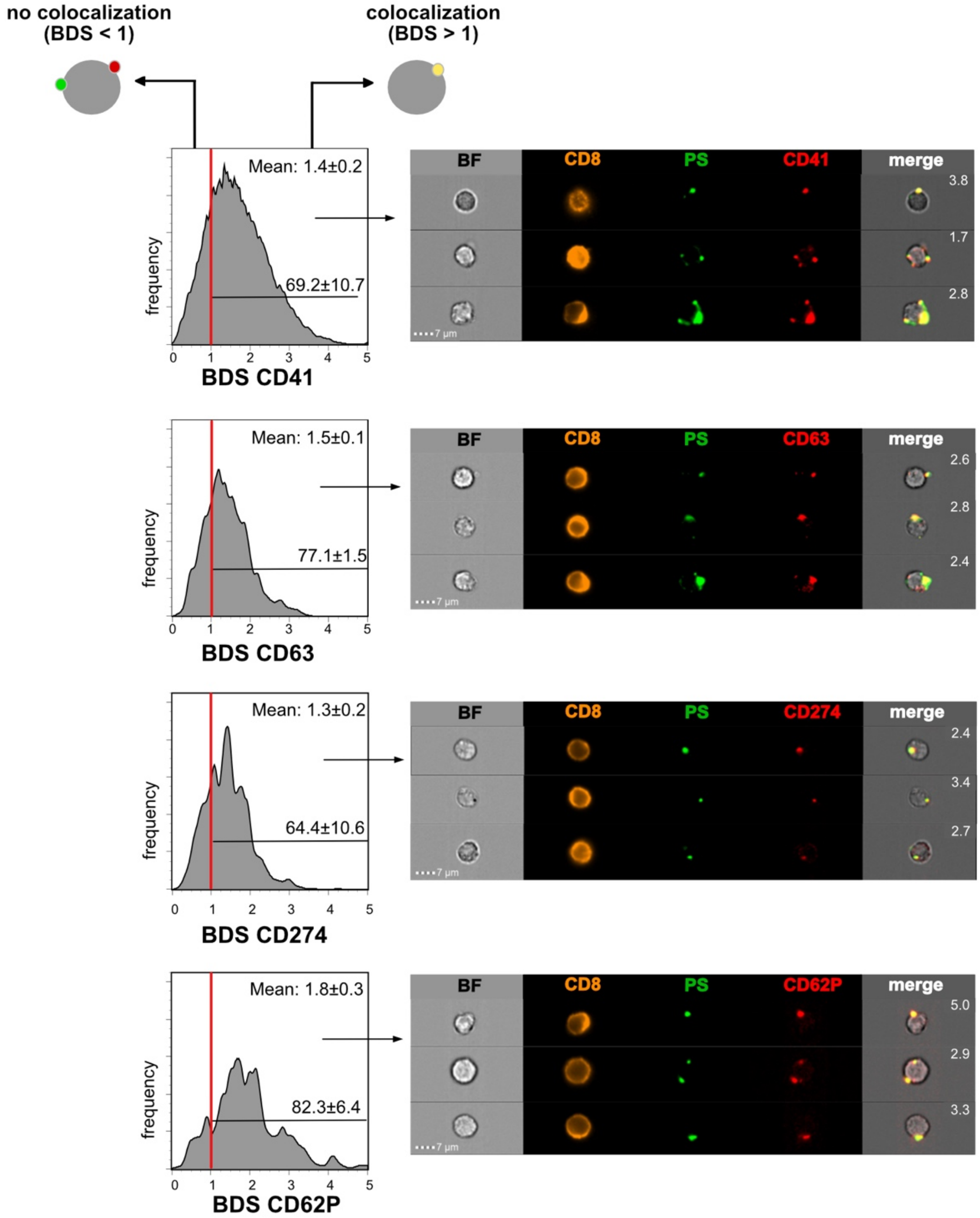


**Suppl. Fig. 5: PS$^+$ EVs colocalize with PMP markers.** Colocalization analysis between PS and PMP-marker staining. To identify PS$^+$ spots the Dilate(Peak(M02, PS, Bright, 5),1) and to identify PMP-marker$^+$ spots the Dilate(Peak(M_marker,marker channel, Bright, 1),1) masks were used. To quantify the degree of colocalization between these masks, bright details similarity scores (BDS) were calculated. (A) Cells with a BDS < 1 did not show any significant colocalization as determined by visual inspection. Cells with a BDS > 1 showed substantial colocalization of PS and the respective platelet marker. BDS scores are shown in the representative example images. Histograms show the BDS scores of PS$^+$EV$^+$ CD8$^+$ cells. The mean BDS score and the percentage of cells showing colocalization (BDS > 1) are indicated within histograms.

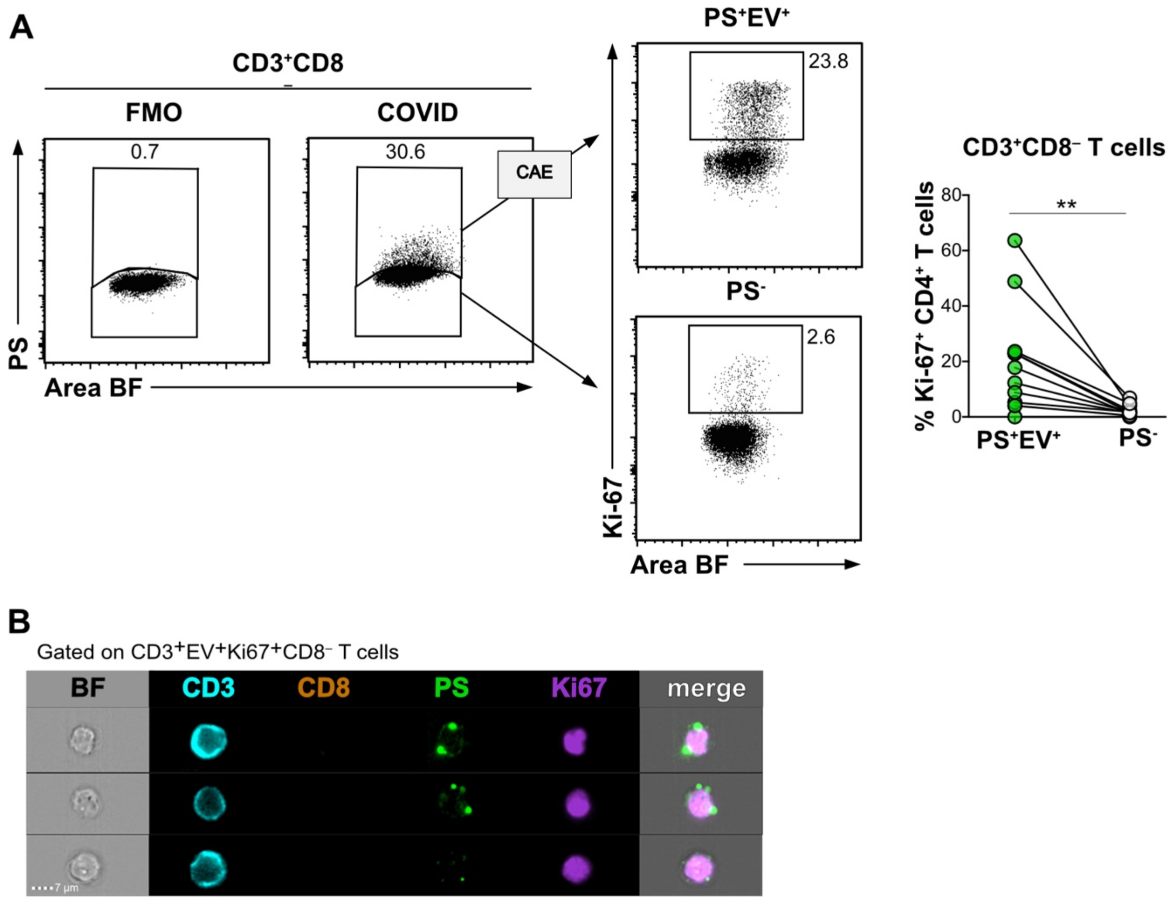


**Suppl. Fig. 6: PS$^{+}$EV associate with proliferating CD4 T cells.** PBMC from COVID patients were examined for proliferation with Ki-67. CD4 T cells were analyzed as shown in Figure Suppl. 1F as CD3$^{+}$CD8$^{–}$ cells. PS$^{+}$CD3$^{+}$CD8$^{–}$ T cells were stained intranuclear for Ki-67. PS$^{+}$ and PS- fractions were classified by IFC. CAE-analysis identified PS$^{+}$EV$^{+}$ live cells (A). PS$^{+}$EV$^{+}$ and PS$^{–}$ CD4$^{+}$ T cells were then analyzed for Ki-67, and data from all patients were plotted on the graph (A). The numbers in the dot plots represent the fraction of cells in the corresponding gate (A). (B) shows an image selection of CD4 T cells with the markers used. Statistical significance was determined by paired Wilcoxon test and is indicated by asterisks (***P ≤ 0.001).

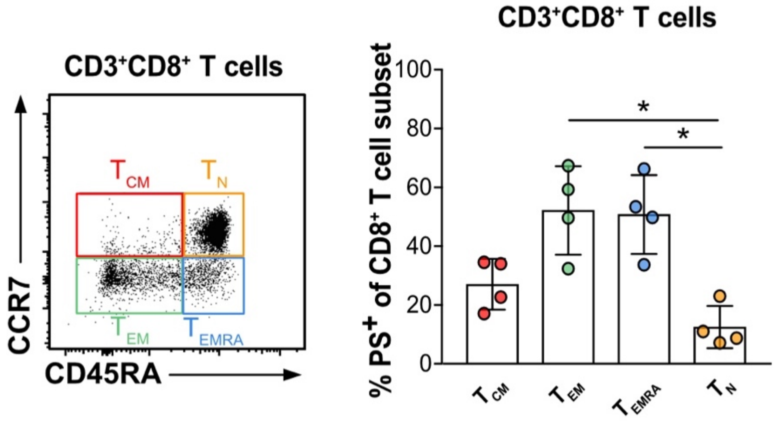


**Suppl. Fig. 7: Analysis of PS$^+$ CD3$^+$CD8$^+$ T cell subsets.** $T_{CM}$ (CCR7$^+$CD45RA$^-$), $T_{EM}$ (CCR7$^-$CD45RA$^-$), $T_{EMRA}$ (CCR7$^-$CD45RA$^+$) and $T_N$ (CCR7$^+$CD45RA$^+$) CD3$^+$CD8$^+$ T cells from COVID patients (n = 4) were examined for PS positivity. PS$^+$ CD8$^+$ T cell subsets were first gated as shown in Figure Suppl. 1 F and then further analyzed based on the expression of CCR7 and CD45RA. The percentage of PS$^+$ cells of the respective CD8$^+$ T cell subset is shown in the graph. Statistical significance was determined by Mann-Whitney test and is indicated by asterisks *P ≤ 0.05
